# Genetic mapping of the early responses to salt stress in *Arabidopsis thaliana*

**DOI:** 10.1101/2020.10.02.324178

**Authors:** Mariam Awlia, Nouf Alshareef, Noha Saber, Arthur Korte, Helena Oakey, Klára Panzarová, Martin Trtílek, Sónia Negrão, Mark Tester, Magdalena M. Julkowska

## Abstract

Salt stress decreases plant growth prior to significant ion accumulation in the shoot. However, the processes underlying this rapid reduction in growth are still unknown. To understand the changes in salt stress responses through time and at multiple physiological levels, examining different plant processes within a single setup is required. Recent advances in phenotyping has allowed the image-based estimation of plant growth, morphology, colour and photosynthetic activity. In this study, we examined the salt stress-induced responses of 191 Arabidopsis accessions from one hour to seven days after treatment using high-throughput phenotyping. Multivariate analyses and machine learning algorithms identified that quantum yield measured in the light-adapted state (Fv′/Fm′) greatly affected growth maintenance in the early phase of salt stress, while maximum quantum yield (QY max) was crucial at a later stage. In addition, our genome-wide association study (GWAS) identified 770 loci that were specific to salt stress, in which two loci associated with QY max and Fv′/Fm′ were selected for validation using T-DNA insertion lines. We characterised an unknown protein kinase found in the QY max locus, which reduced photosynthetic efficiency and growth maintenance under salt stress. Understanding the molecular context of the identified candidate genes will provide valuable insights into the early plant responses to salt stress. Furthermore, our work incorporates high-throughput phenotyping, multivariate analyses and GWAS, uncovering details of temporal stress responses, while identifying associations across different traits and time points, which likely constitute the genetic components of salinity tolerance.

## Introduction

High soil salinity is threatening global food security in the midst of rapid population growth and climate change (Godfray et al. 2010; Alexandratos and Bruinsma 2012; Qadir et al. 2014), which has been historically reported to be caused by flooding, over-irrigation, silting and the rising water table (Singh et al. 2010; Daliakopoulos et al. 2016; Shahid, Zaman, and Heng 2018). Soil salinisation can limit crop productivity in arid, semi-arid and coastal areas by interfering with nutrient and water uptake, resulting in an annual yield loss of ∼27.3 billion USD (Qadir et al. 2014; Butcher et al. 2016). Plants respond to increased soil salinity in two phases, reflecting an early osmotic phase and a late ionic phase (Munns 2002; Munns and Tester 2008). During the later ionic phase, which has been more widely studied, salt ions accumulate to toxic levels in the shoot tissue, leading to premature leaf senescence and decreased plant productivity (Kobayashi et al., 2017; Shamaya et al., 2017; Wu et al., 2018). However, rapid growth reduction has been consistently observed prior to ion accumulation in the shoot across different plant species (Rajendran et al., 2009; Adem et al. 2014; Tilbrook et al. 2017). This early phase of salt stress response is referred to as the osmotic phase (Munns and Tester, 2008), and the underlying mechanisms were proposed to contribute to shoot-ion independent tolerance (Roy et al., 2014). During the osmotic phase, the reduction in growth is due to a decrease in cell cycle activity (West et al., 2004), cell expansion (Niu et al. 1996; Neves-Piestun and Bernstein 2005), leaf emergence (Zeng and Shannon 2000; James et al. 2002; Fricke et al. 2006; Luo et al. 2017) and cell wall rigidity (Byrt et al. 2018; Feng et al. 2018). Hence, mechanisms underlying ion sensing, cell cycle regulation and stomatal conductance are hypothesised to contribute to the reduction in growth during the osmotic phase (Muranaka et al., 2002; West, Inzé, and Beemster 2004; Zhu 2016; W. Gao et al. 2018). However, the genetic constituents of the maintenance of growth during the osmotic phase of salt stress are still largely unknown.

By performing regular growth measurements after salt stress exposure for an extended period of time, one can discriminate between the early and late responses to salt stress (Roy et al., 2014). In addition, salt stress has been reported to increase the leaf temperature (Rahnama et al. 2010; Campos et al. 2016; Hu et al., 2017), decrease the photosynthetic efficiency (Pan et al. 2016; Percey et al. 2016), reduce the growth rate (West et al., 2004; Zeng and Shannon 2000) and change the leaf colour (Awlia et al. 2016). However, to the best of our knowledge, the contribution of each of these traits to the overall plant performance under salt stress has not yet been quantified. This would require that all the traits be recorded in a single experiment in a high-throughput manner. Fortunately, the advances in non-destructive high-throughput phenotyping have allowed the quantification of a variety of traits through time (Humplík et al. 2015; Fahlgren et al., 2015; Tardieu et al. 2017), revealing the dynamic relationships between the traits (York 2019) and quantifying even the subtle changes induced by stress exposure (Awlia et al. 2016). The automated nature of high-throughput phenotyping also allows the screening of large populations of plants within a relatively short amount of time, making it suitable for forward genetic studies. These studies typically require screening large populations of mutagenised plants, bi-parental crosses or natural diversity panels. Large-scale sequencing projects, such as the 1,135 Arabidopsis genome project (1001 Genomes Consortium), provided the scientific community with high-quality and high-density single nucleotide polymorphism (SNP) markers, enabling high-resolution mapping by genome wide association studies (GWAS). While GWAS have been successfully deployed for the identification of genetic components underlying ion accumulation (Baxter et al. 2010), root growth (Y. Kobayashi et al. 2016; Julkowska et al. 2017; Deolu-Ajayi et al. 2019) and biomass accumulation (Julkowska et al. 2016) under salt stress, each of these approaches only focused on a specific set of phenotypes within one or a few time points.

In this study, we phenotyped 191 Arabidopsis accessions using a high-throughput phenotyping platform and an optimised salt-application protocol that was developed for soil-grown plants (Awlia et al. 2016). Plant growth, colour and photosynthetic efficiency were measured daily from one hour to seven days after salt treatment. Our comprehensive multivariate analysis revealed the dynamics of the salt-induced changes across the measured traits and the relative contribution of each trait to the overall plant productivity. The collected 438 phenotypes, representing 50 unique traits scored over eight time points under control and salt stress conditions. In addition, the growth rate was estimated across two time intervals. All measured and derived traits were used as inputs for the single and multi-trait GWAS models to identify the salt stress-specific associations. From the identified 770 loci, we focused on the genomic regions associated with photosynthetic efficiency in the form of quantum yield (QY max and Fv′/Fm′), as these traits were strongly correlated with growth maintenance under salt stress. This correlation was determined by a machine learning approach. Through phenotyping the available T-DNA insertion lines of the genes within the Fv′/Fm′ locus, we identified an uncharacterised protein kinase that seems to play an important role in regulating photosynthetic efficiency and plant productivity under salt stress.

## Materials & Methods

### Plant material and growth conditions

In this study, 191 accessions of the HapMap population (Alonso-Blanco *et al*., 2016) were used (seeds provided by Prof. Christa Testerink, Universiteit van Amsterdam), in which the accessions that were reported to have a dwarf phenotype or low seed germination were excluded (**Table S1**). The seeds were soaked in Milli-Q water and stratified for three days at 4 °C in the dark. The seeds were sown into pots (70 x 70 x 65 mm, Pöppelmann TEKU, GmbH, Germany) containing 60 g of freshly sieved soil (Substrate 2, Klasmann-Deilmann GmbH, Germany) and watered to full soil-water holding capacity using tap water. All plants were grown in a climate-controlled growth chamber (FS_WI, Photon Systems Instruments, Czech Republic) with cool-white light emitting diode (LED) in a 12/12 h 22°C/20°C light/dark cycle with a relative humidity of 55% and a white-light photon irradiance of 150 µmol m^-2^ s^-1^ (**Figure S1 A**).

### Salt stress treatment and high-throughput phenotyping of the HapMap population

The soil water-holding capacity was determined by drying ten pots of sieved soil at 80 °C for three days, which were weighed when fully saturated with water. By using 60% of that capacity, the soil-water content was controlled prior to salt treatment (**Figure S1 A**). At 14 days after germination (DAG), the seedlings were randomly allocated to different positions within the trays (5 x 4 pots per tray) according to a split-plot block design (**Figure S2**) in the PlantScreen^TM^ conveyor system (PSI, Czech Republic). The pairs of salt and control-treated trays were designed to contain the same genotypes at the same positions (**Figure S2**) and were kept adjacent to one another during the experiment. The replicates of each accession were distributed across the different tray pairs, in which 4 to 6 biological replicates were included per accession and condition. The plants that died during the experiment or did not exhibit any change in growth were excluded from the analysis, reducing the number of replicates for some accessions to be as low as 1, which were also excluded. Thus, the accessions that did not include a representative plant under the conditions of both control and salt stress were excluded from the analysis. Automatic weighing and watering were performed every other day to maintain the reference weight that corresponded to 60% of the soil-water holding capacity. At the 10-leaf stage (24 DAG), the pots were saturated for one hour in tap water or a 250 mM NaCl solution, resulting in a final concentration of 100 mM NaCl based on the 60% soil-water content, which corresponded to a moderate level of salt stress (Awlia *et al*., 2016). Pots were then placed in the PlantScreen^TM^ system for [red:green:blue] (RGB) (**Figure S1 B**) and chlorophyll fluorescence imaging after one hour of treatment and through seven consecutive days, and then returned to the same positions inside the growth chamber. A single round of measuring consisted of an initial 15-min dark-adaptation period inside the acclimation chamber, followed by chlorophyll fluorescence and RGB imaging. Light conditions, plant position and camera settings were fixed throughout the measurements. In addition, the rate of photosynthesis was quantified at different photon irradiances in the form of light steady states (Lss) consisting of Lss1 to Lss4 that corresponded to 95, 210, 320 and 440 µmol m^-2^ s^-1^, respectively, using the light curve protocol as described by <u>Henley (1993)</u>. During the phenotyping period, the plants were not re-watered to avoid diluting the NaCl concentration in the soil. On the final day of imaging, the soil-water content was measured to be between 70-80%, indicating that the plants had adequate soil moisture during the experiment.

### Data analysis

The raw phenotypic data of the measured Arabidopsis HapMap accessions in this study can be found at https://doi.org/10.5281/zenodo.3740415, while the R notebook summarising the analysis steps is available at https://doi.org/10.6084/m9.figshare.12173382.v1. Furthermore, the change in rosette area over time, as determined by the RGB pixel counts, was used to estimate the growth rate per accession. A linear function (y = A + B * time (days), where A is the starting rosette area and B is the growth rate in mm^2^ per day) was fitted over the two time intervals of 0 to 3 and 4 to 7 days after treatment using a linear regression model and the lm function in R v4.0.0 (R Core Team, 2020), which was primarily used for data analysis. The relative performance of each accession under salt stress was estimated by dividing its average growth rates (GR) under salt stress by the average under control conditions (GR _(av salt)_/GR _(av control)_) per interval, reflecting a salt tolerance index for intervals 1 and 2. The correlation analysis was performed using the corrplot R package (v0.84) (Wei and Simko, 2017) on the average value per accession and condition, which was calculated using the doBy package (v4.6.6) (Højsgaard *et al*., 2016). The machine learning model was executed using the caret (v6.0-86), xgboost (v1.0.0.2) and mlbench (v2.1-1) R packages (Kuhn, 2015; Chen *et al*., 2018; Leisch and Dimitriadou, 2010), while the graphs were produced using the ggplot2 (v3.3.0), ggpubr (v0.3.0) and cowplot (v1.0.0) R packages (Wickham, 2016; Wilke, 2016).

### Spatial correction and genome wide association mapping

Spatial correction was applied on the phenotypic data using the ASReml R package (v3.5.0) (Butler *et al*., 2009) to account for the possible spatial effects of position, microenvironments and shading (**Figure S2 A**). The spatial analysis was based on a split-plot block design that accounted for the pot position (sub-plot), tray position (main-plot), the four pairs of trays on a bench (bench-row) and the position of the tray within the growth chamber, where the replicates were randomly placed among the trays. Spatial correction of the rosette area on the final day of imaging was used as a proof of concept using the number of bench-rows within the split-plot block design (**Figure S2 B-C**). In addition, the changes in each measured trait in response to salt stress were evaluated using a one-way analysis of variance (ANOVA) using the grand average of the population across both control and salt stress conditions. The spatially corrected data has been made available at https://doi.org/10.5281/zenodo.3740429, which was used as an input for the GWAS.

Genome wide association mapping was run in R using the EMMAX and the ASReml R package (v3.5.0) (Butler *et al*., 2009) using single and multi-trait mixed models (MTMM) that study the within-trait and between-trait variance by testing the genotypic, environmental or genotype by environment effects on the association strength. The GWAS script included terms for correcting the population structure and the kinship matrix (Korte *et al*., 2012). To determine the population structure, principal component analysis (PCA) was applied using the factoextra R package (v1.0.7) (Kassambara and Mundt, 2017). While the GWAS was run using 4,285,827 SNPs, we only considered SNPs with a minor allele frequency above 0.05 to exclude rare alleles. Therefore, the Bonferroni threshold was implemented using the –log_10_(*p*-value / number of SNPs) for SNPs with a minor allele frequency above 0.05. This corresponded to a Bonferroni threshold of - log_10_(0.05/1,753,576) = 7.55. Moreover, the MTMM can improve the odds of detecting significant SNPs that are specific to genotype, environment or both, which reflect the environment-independent, environment-specific or full test analyses, respectively. Therefore, we focused on the significant associations that were detected using the environment-specific and full tests, representing salt-specific associations. However, model convergence is a common problem in MTMM analyses, especially when the within-genotype variance is high. In this study, successful convergence of the environment interaction model was only observed in 29 out of the 219 pairs of traits that were measured under salt and control conditions to assess the genetic association across treatments. The quantile-quantile (Q-Q) plots of the observed and expected *p*-values of the GWAS results were generated using the qqman (v0.1.4) and qvalue (v2.20.0) R packages (Turner, 2011; Dabney *et al*., 2010). This was performed to indicate the *p*-value inflation and examine model fitting, which was based on curve skewness (**Figure S5, Figure S7, Figure S10**). The GWAS output files can be found at https://doi.org/10.5281/zenodo.3740443, while the R script used to generate the Manhattan and QQ plots can be found at https://rpubs.com/mjulkowska/ArabidopsisEarlySaltManhattanAndQQ.

As the average linkage disequilibrium in Arabidopsis has been estimated to be 10 kbp (Nordborg *et al*., 2002), we considered the 10 kbp upstream and downstream of the significantly-associated SNP as one locus. Hence, the neighbouring associated SNPs within that region were considered part of one locus, which was screened for open reading frames (ORFs) that encoded putative candidate genes (**Figure 5**, **Figure 7**). The significant associations of SNPs that were repeatedly observed across different traits and time points were identified and filtered. Next, the 684 most promising loci were ranked by the -log_10_(*p*-value) as well as the total number of associated SNPs and traits. The significant SNPs within the two most promising loci, which represented the light- and dark-adapted quantum yield (F_v_′/F_m_′ and QY max, respectively), were investigated using the Arabidopsis thaliana 1001 Genome Browser (Salk Institute) and The Arabidopsis Information Resource (TAIR). Their expression patterns were analysed using the Arabidopsis eFP Browser (Kilian *et al*., 2007; Winter *et al*., 2007; Bassel *et al*., 2008) under control and salt stress conditions within the root and shoot using the natural variation therein. In addition, the predicted subcellular localisation was explored using the subcellular localisation database for Arabidopsis proteins (SUBA) (Hooper *et al*., 2017).

### Functional validation of the identified loci using T-DNA insertion lines

The T-DNA insertion lines of the selected candidate genes (**Table S4**) were ordered from the Nottingham Arabidopsis Stock Centre (NASC) and validated using the primers designed by The SALK T-DNA Express (**Table S5**). The DNA was isolated using the Edwards DNA extraction protocol (Edwards *et al*., 1991), where Col-0 served as a negative control for the PCR reactions. The homozygous lines were propagated in the same batch with the Col-0 background line, in which the harvested seeds were used for phenotyping under control and salt stress conditions. Furthermore, the effect of the T-DNA insertion on the gene expression was tested for soil-grown plants at 28 DAG using the leaf tissue of three biological replicates. The RNA was extracted using TRIzol (dx.doi.org/10.17504/protocols.io.pbndime) and treated with DNAse using Invitrogen™ DNA-free™ DNA Removal Kit (AM1906, Invitrogen Corp., Thermo Fisher Scientific, USA). In addition, the cDNA was synthesised using the Bio-Rad iScript™ cDNA Synthesis Kit (25 x 20 µl reactions #1708890, Bio-Rad Laboratories Inc., USA), while the qPCR reaction was performed using the Applied Biosystems PowerUp™ SYBR™ Green Master Mix (A25741, Applied Biosystems, Thermo Fisher Scientific, USA). The Arabidopsis actin (AtActin) primers were used to normalise the expression for the primers used in each gene expression (**Table S6**). Finally, the expression of the target genes within each T-DNA insertion line was examined using three biological replicates, where the expression levels were calculated using the delta-Ct method relative to the AtActin values.

The Col-0 and the homozygous T-DNA insertion lines (**Table S5**) were grown under the same conditions as was used in the high-throughput phenotyping setup (12/12 h 22°C/20°C light/dark cycle with a relative humidity of 55% and a white-light photon irradiance of 150 µmol m^-2^ s^-1^). Salt stress was applied at 14 DAG, where each pot was placed in tap water or a 250 mM NaCl solution for 10 mins before being placed in the phenotyping system PlantScreen^TM^ (KAUST, Saudi Arabia) for RGB and chlorophyll fluorescence imaging as well as regular weight measurements, which was performed through eleven consecutive days after salt treatment. The light curve chlorophyll fluorescence protocol was applied during 60 s intervals of cool-white actinic light at photon irradiances of 95, 210, 320, 440, 555 and 670 µmol m^-2^ s^-1^ corresponding to Lss1 to Lss6, respectively. In addition, each T-DNA insertion line was represented by 12 biological replicates per treatment, while Col-0 was placed in each tray as a reference. During the phenotyping period, the plants were not re-watered to avoid diluting the NaCl concentration in the soil. The raw phenotypic data can be accessed at https://doi.org/10.5281/zenodo.3762244, while the R-analysis pipeline is available at https://rpubs.com/mjulkowska/Arabidopsis_PSI_TDNA_pipeline_01.

## Results

### The natural variation in salt stress responses of Arabidopsis

A total of 191 Arabidopsis accessions of the HapMap population were examined in their early responses to salt stress (**Table S1**) using the high-throughput phenotyping platform, PlantScreen^TM^ (**Figure S1 A**). The changes in rosette size, morphology and chlorophyll fluorescence were captured daily from one hour to seven days after salt treatment of 100 mM NaCl. A number of accessions treated with salt stress showed significantly smaller rosettes relative to the control-treated plants starting from one hour after salt treatment (**Figure 1 A**), and the significant differences between the salt and control-treated plants were found to increase over time. In addition, analysis of plant morphology (**Figure S1 B**) revealed that under salt stress plants developed more compact rosettes (**Figure 1 B**) and less slender leaves (**Figure S3 A**) compared to control-treated plants. Furthermore, significant changes in rosette morphology were observed three days after treatment (**Figure 1 B**), classifying them as relatively late salt stress responses. We also extended the analysis of the RGB images by segmenting the Arabidopsis rosette into nine representative green hues and calculated the relative hue abundance throughout the experiment as a fraction of the rosette area (**Figure S4**). The significant difference in hue abundance between plants grown under control and salt stress conditions was apparent as early as one hour after salt treatment for hues 1 and 5, which represented pale-green and moss-green, respectively (**Figure S4**). One day after salt treatment, the relative abundance of darker hues (hues 4 and 6) increased, while the abundance of lighter hues (hue 3) decreased. The most abundant hue (hue 2) was only slightly decreased in salt-treated plants six days after treatment (**Figure S4**). Hence, the changes in rosette colouring can be classified as being both early and late salt stress responses, depending on the specific hue and its overall abundance.

**Figure 1.**
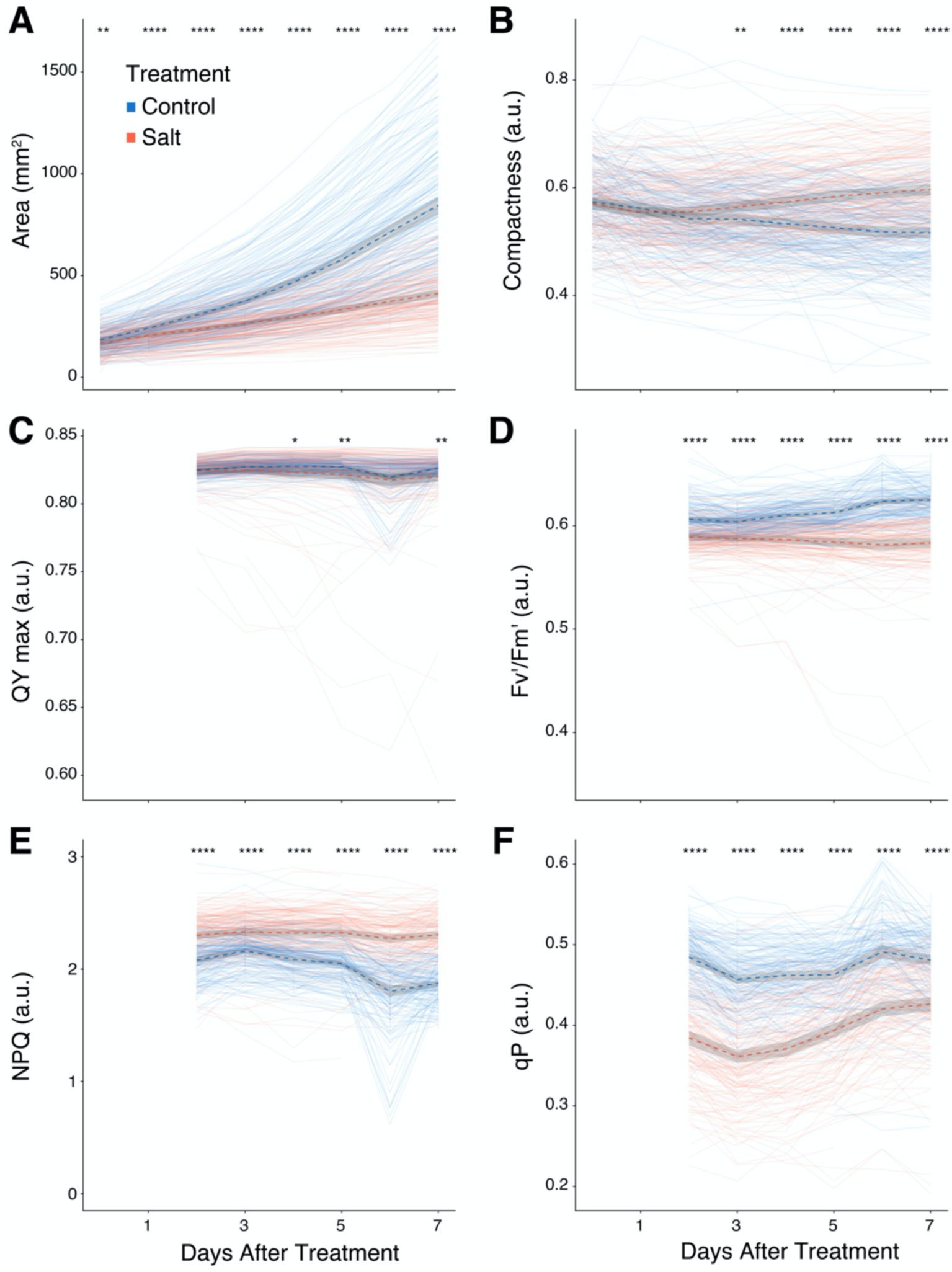
Salt stress can affect plant size, morphology and photosynthetic efficiency. The salt stress responses of three-week-old soil-grown plants that were exposed to a final concentration of 100 mM NaCl, were examined as described by (Awlia *et al*., 2016). **(A)** Rosette area (mm^2^) and **(B)** compactness (ratio of area to surface of convex hull) were calculated based on the pixels captured by the RGB camera immediately after salt treatment and seven days thereafter. **(C)** Maximum quantum yield was only captured two days after salt treatment (due to technical difficulties) in the dark-adapted (Fv/Fm, referred to as QY max) and **(D)** the light-adapted (F_v_′/F_m_′) states using the highest photon irradiance of 440 μmol m^-2^ s^-1^ with the light curve protocol of chlorophyll fluorescence imaging. **(E)** Non-photochemical quenching (NPQ=[F_m_– F_m_′]/(F_m_′)) and **(F)** the photochemical quenching coefficient (qP=[F_m_′–F_t_]/[F_m_′–F_o_′]) were determined by estimating the maximum fluorescence in the dark (F_m_) and light-adapted (F_m_′) states as well as the instantaneous fluorescence (F_t_) and the minimal fluorescence (F_o_′) in the light-adapted state. Each line represents the trajectory per accession through time, where blue and red lines indicate plants grown under control or salt stress conditions, respectively. The dashed lines represent the mean of the population per condition and trait, while the grey band represents the confidence interval. The asterisks above the graphs indicate significant differences between control and salt-treated plants within the entire population, where *, **, *** and **** indicate *p*-values < 0.05, 0.01, 0.001 and 0.0001, respectively, as determined by one-way ANOVA.

To examine the salt-induced changes in photosynthetic efficiency, chlorophyll fluorescence imaging was performed using the light curve protocol <u>(Henley 1993)</u>, which was only measured two days after salt treatment due to technical difficulties. The maximum quantum yield (QY max), calculated as a ratio of variable (F_v_) to maximum fluorescence (F_m_) in the dark-adapted state, was only slightly reduced by salt stress and its effect was not significant until four days after treatment (**Figure 1 C**). However, a small number of accessions exhibited a severe decrease in QY max, from the typical value of ∼0.83 in the absence of stress (Murchie and Lawson, 2013) to 0.65 after salt treatment (**Figure 1 C**). The ratio of light-adapted variable (F_v_′) and maximum fluorescence (Fm′), which reflect the light-adapted quantum yield (F_v_′/F_m_′), also showed significant decreases under salt stress. However, the salt-induced changes were observed as early as two days after salt treatment (**Figure 1 D**). Furthermore, non-photochemical quenching (NPQ=[F_m_–F_m_′]/(F_m_′)), which is antagonistic to photosynthetic activity and chlorophyll fluorescence, was observed to increase in response to salt stress (**Figure 1 E**), while the photochemical quenching coefficient (qP=[F_m_′–F_t_]/[F_m_′–F_o_′]) decreased (**Figure 1 F**), where both NPQ and qP showed significant changes as early as two days after salt treatment. Moreover, salt stress induced rapid changes in the minimal (F_o_′) and maximal (F_m_′) chlorophyll fluorescence parameters (**Figure S3 B-D**), where a significant decrease was observed two days after salt treatment (**Figure S3 C-D**). In contrast, the instantaneous chlorophyll fluorescence (F_t_) parameter was not significantly decreased until five days after salt treatment (**Figure S3 B**). In summary, the dark-adapted chlorophyll fluorescence parameters showed relatively late responses to salt stress, while the light-adapted chlorophyll fluorescence parameters as well as the photochemical and non-photochemical quenching parameters were more affected by salt stress at earlier time points, except in the case of Ft. Taken together, the salt-induced changes that were observed during the seven days after salt treatment consisting of rosette area, greenness hues and the light-adapted chlorophyll fluorescence parameters displayed rapid changes in response to salt stress, while rosette morphology, the dark-adapted quantum yield and the instantaneous fluorescence displayed later changes after salt treatment.

### The early and late responses to salt stress were not correlated

Changes in rosette area, across two time intervals of 0 to 3 and 4 to 7 days after salt treatment, were investigated, while corresponding to the early and late responses to salt stress, respectively. Furthermore, a linear function, representing the change in rosette area through time, was fitted per accession and condition (**Figure S5**). In general, the growth rates of control-treated plants were higher in the second interval relative to the first interval (**Figure 2 A**), while the growth rates of salt-treated plants were lower than that of the control-treated plants. The difference between the conditions increased during the second interval, suggesting an accumulative effect of salt stress over time (**Figure 2 A**). Moreover, as the growth rate of control-treated plants increased in the second interval, we also observed greater natural variation in the growth rate between the accessions.

**Figure 2.**
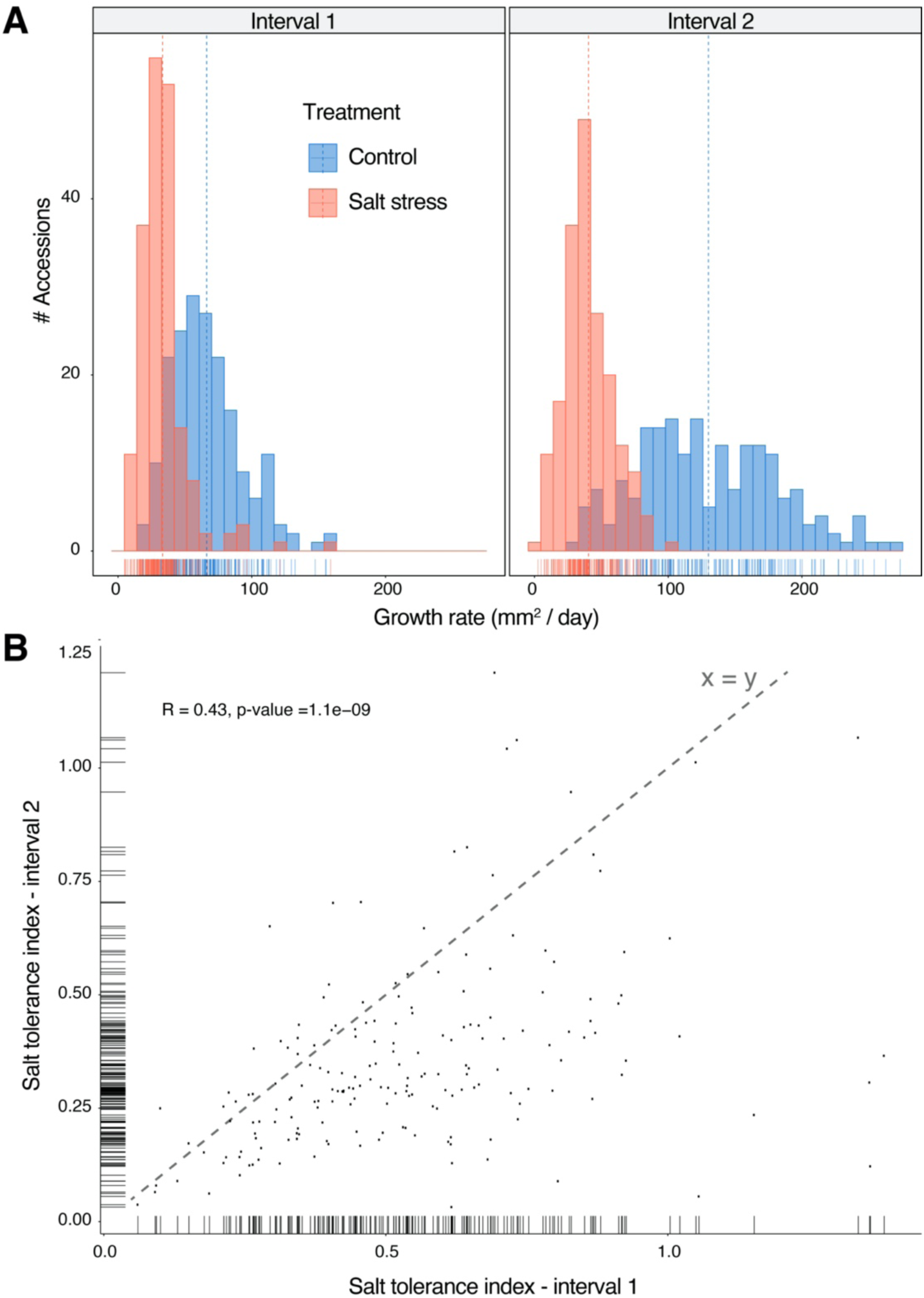
Growth maintenance during the early and late stages of salt stress is not correlated. **(A)** The growth rate (mm^2^/day) of each accession was estimated by fitting a linear function to the change in rosette area across the two time intervals of 0 to 3 and 4 to 7 days after treatment. The histogram shows the distribution of growth rates during intervals 1 and 2 (left and right panel, respectively) across the control (blue) or salt-treated (red) accessions. **(B)** The salt tolerance index (STI) was calculated per interval by dividing each accession’s growth rate that was measured under salt stress by the growth rate measured under control conditions. The scatter plot represents the correlation between STI calculated for intervals 1 and 2, while the marginal rugs on the x- and y-axis represent the distribution of STI across each interval. The correlation coefficient (R) and *p*-value can be found in the upper left-hand corner, while the dashed diagonal line represents the position of the points when STI-interval 1 is equal to STI-interval 2 (y = x).

To estimate the performance of each accession per interval, we calculated the salt tolerance index per interval by dividing the growth rate under salt stress by the growth rate under control conditions. Salt stress clearly caused a more pronounced growth reduction in the second interval, as the second interval index values were lower than that of the first interval (**Figure 2** **B**). Next, the correlation across the two intervals of the salt tolerance index was examined (**Figure 2 B**). Although the *p*-value of the correlation coefficient (R) indicated a significant correlation, the correlation itself was weak (R < 0.5), where a substantial number of accessions varied in their performance across the first and second intervals (**Figure 2 B**). This suggested that the mechanisms contributing to the maintenance of plant growth across the two intervals were distinct and might involve different genetic components and molecular processes that are unique to the early and late responses to salt stress. To identify the genetic components contributing to the growth rate at each interval and under control and salt stress conditions, single-trait GWAS was implemented, in which no significant associations were identified (**Figure S6, A-B and D-E**). One significantly-associated SNP was found in the first interval using the salt tolerance index (chromosome 2, position 5393189) (**Figure S6 C**) and nine in the second interval, with four of those SNPs were found on chromosome 5 between positions 6221844 and 6247148 (**Figure S6 F**). Since the associations with the salt tolerance index were not particularly robust, we chose to investigate associations with other measured traits.

### Identifying traits contributing to growth maintenance under salt stress

Our phenotyping dataset included 50 unique traits that were measured across multiple days and two conditions, from which we derived growth rates and stress indices (**Table S2**). To gain insight into the interdependent relationships between traits as well as their relationship to growth maintenance, we performed multivariate analyses. By studying the relationships between the measured traits under control and salt stress conditions (**Figure 3 A - B**), we observed strong correlations between rosette area, perimeter, rosette compactness, roundness and the directly-measured chlorophyll fluorescence traits (Fm, Fo, Ft and Fv). Moreover, the correlations between most chlorophyll fluorescence traits and non-photochemical quenching (NPQ) were negative, while the correlation coefficient between maximum quantum yield (QY max) and quantum yield in the light-adapted conditions (Fv′/Fm′) increased under salt stress (**Figure 3 A-B**). Interestingly, the correlation between QY max and rosette area was only significant under salt stress (**Figure 3 B**). This could be due to a small number of accessions that showed a compromised QY max under salt stress, which also developed smaller rosettes (**Figure 3 C**). This indicates the importance of maintaining QY max within the range of 0.80 and 0.83 (Murchie and Lawson, 2013).

**Figure 3.**
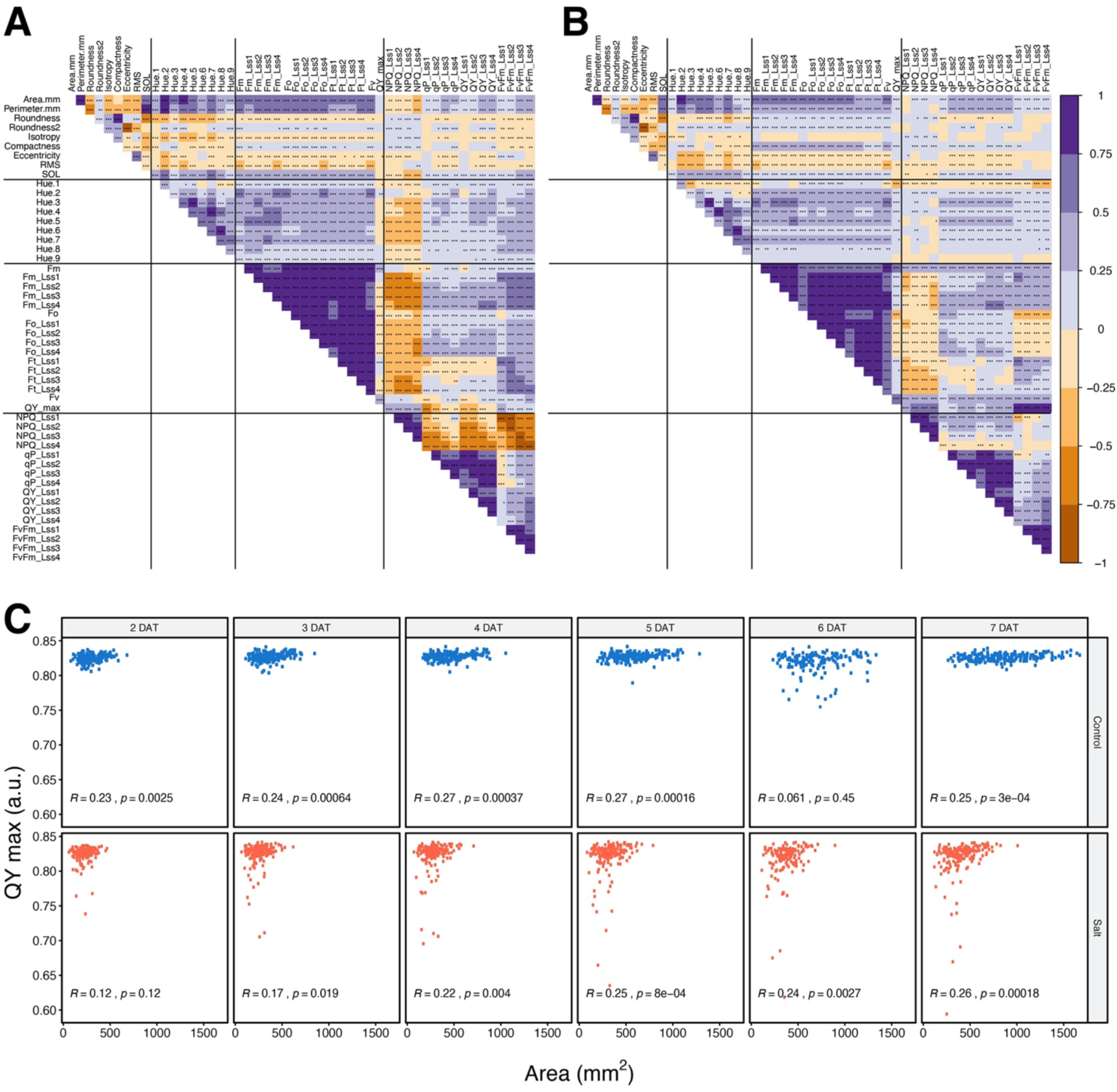
Correlation analysis reveals the salt-specific relationship between QY max and rosette area. The Pearson correlation test was performed on the traits measured under **(A)** control and **(B)** salt stress conditions across all time points and irradiances, where Lss1-4 represents the chlorophyll fluorescence measured under photon irradiances of 95, 210, 320 and 440 μmol m^-2^ s^-1^, respectively. Positive and negative correlations are shown as purple and orange hues, respectively, while the asterisks indicate significant correlations within the entire population, in which *, **, *** and **** indicate *p*-values < 0.05, 0.01, 0.001 and 0.0001, respectively. **(C)** The correlation between rosette area and maximum quantum yield (QY max), which reflects the ratio of variable fluorescence (Fv) to maximum fluorescence (Fm) in the dark-adapted state, was examined under control and salt stress conditions using scatter plots from 2 to 7 days after treatment (DAT). The correlation coefficient (R) and the *p*-value of each correlation can be found in the bottom left-hand corner.

To identify other important traits that could contribute to rosette area under control and salt stress conditions, we developed a machine learning model using gradient-boosted trees. We used 50% of the 1,102 data points to act as a training dataset, while limiting the tree depth to six levels to reduce the risk of memorisation. Subsequently, the remaining 50% of the data was used to test the model’s predictions (**Figure 4 A**). We found that the model was able to predict rosette area with high accuracy, as reflected by the root mean square error (RMSE) of 18.6 and 17.1 mm^2^ in control and salt-treated plants, respectively (**Figure 4 A**). We then examined the trait ranking in each model, where perimeter and greenness hue 2 were ranked the highest (**Figure 4 B**). Since perimeter was highly correlated with rosette area (**Figure 3 A-B**), it was not surprising that it ranked high in predicting rosette area. Hue 2, on the other hand, was the most dominant hue in rosette area under both control and salt stress conditions (**Figure S4**). In the second iteration, we excluded perimeter and all the hue traits from the dataset to construct a new model (**Figure 4 C**), in which the prediction of rosette area was slightly less accurate, with RMSE values of 23.7 and 23.8 mm^2^ in control and salt-treated plants, respectively. The highly ranked traits included slenderness of leaves (SOL) and eccentricity, reflecting rosette morphology, across both control and salt-treated plants (**Figure 4 D**). The chlorophyll fluorescence parameters measured in the dark-adapted state (Fv, QY max) had high contributions to rosette area in salt-treated plants, while the parameters measured under the highest photon irradiance (Fm_Lss4 and Fo_Lss4) showed the highest contribution to rosette area in control-treated plants.

**Figure 4.**
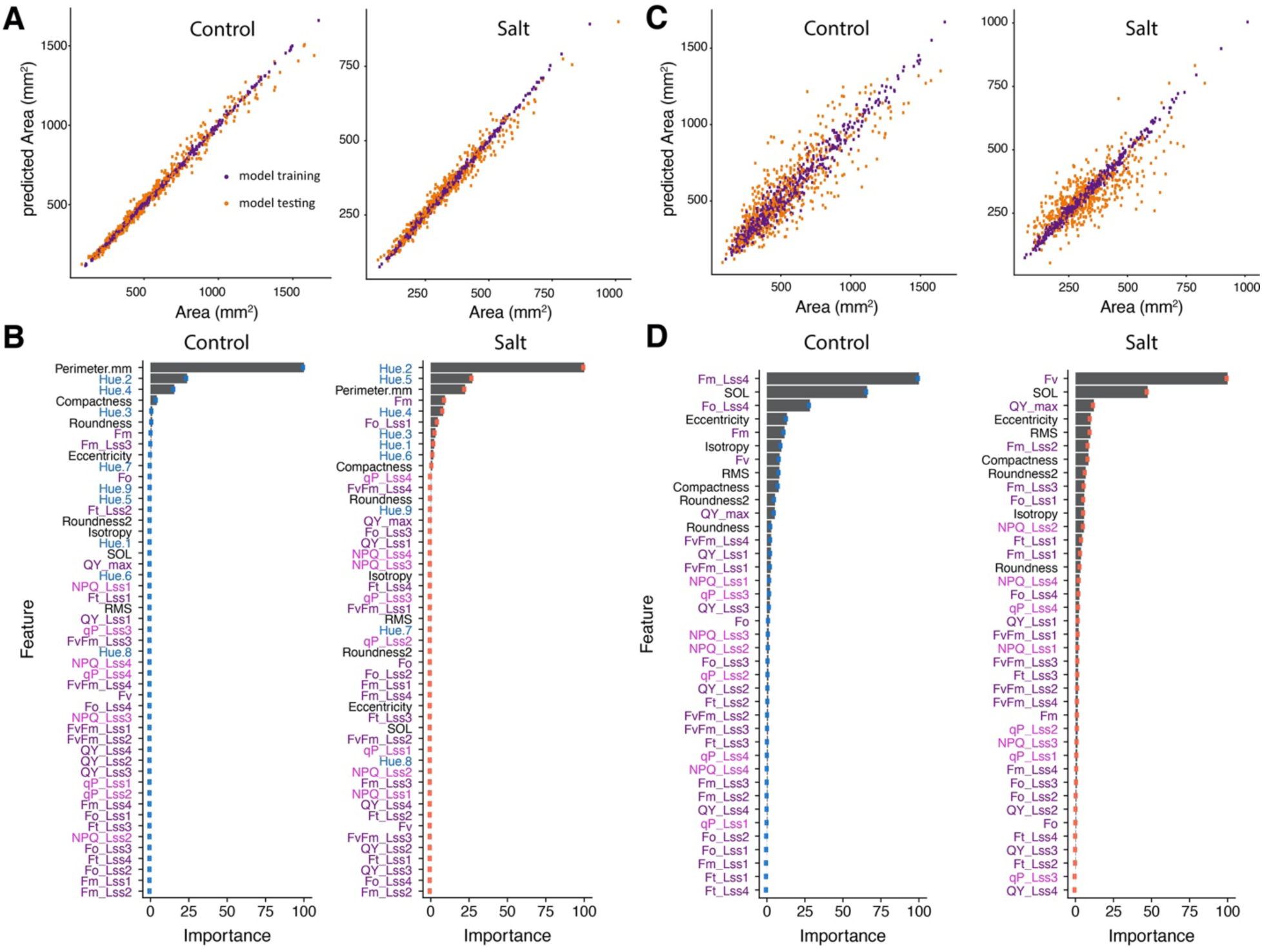
Machine learning models can identify the traits that contribute the most to rosette area. **(A)** The gradient-boosted tree model was developed using 50% of the phenotypic data to predict rosette area (mm^2^) based on all the measured traits under control and salt stress conditions. Data points used for model training (purple) or testing (orange) are shown. **(B)** Feature-scaling analysis, which normalises the independent features present in the data in a fixed range, ranked the importance of the traits that were included in the machine learning model in (A) that predicted rosette area. Traits related to rosette morphology (black), rosette colouring (blue), photosynthetic efficiency (purple) and photochemical or non-photochemical quenching (pink) are highlighted. **(C)** A second gradient-boosted tree model was developed using 50% of the data to predict rosette area based on all the measured traits with the exception of rosette perimeter and the greenness hues. **(D)** Feature-scaling analysis ranked the traits that were included in the machine learning model in (C) that predicted rosette area.

**Figure 5.**
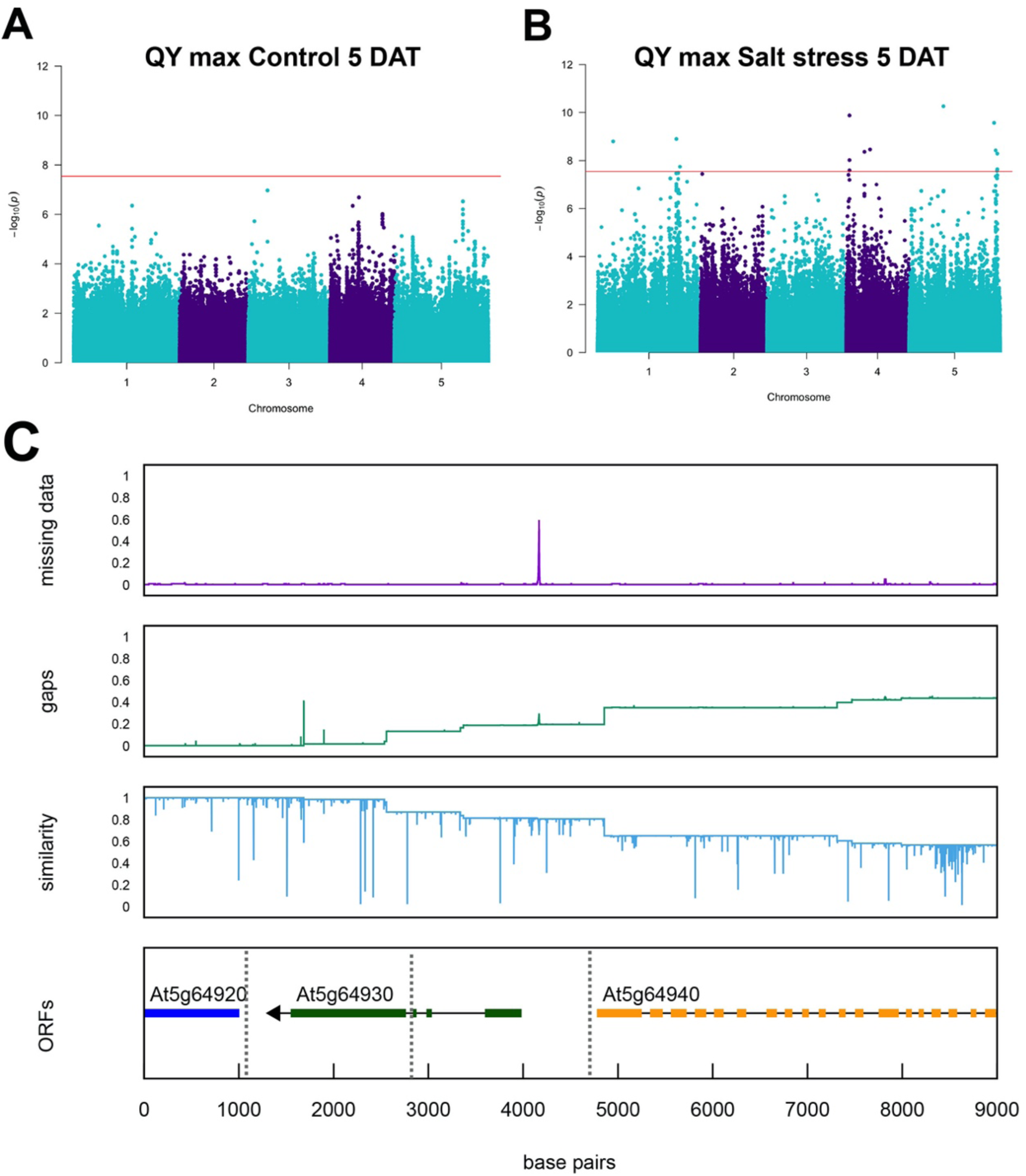
Identifying the genetic loci underlying maximum quantum yield under salt stress. Single-trait genome-wide association studies (GWAS) was conducted on the maximum quantum yield of photosystem II photochemistry in the dark-adapted state (QY max = Fv/Fm) that was measured under **(A)** control and **(B)** salt stress conditions five days after treatment (DAT). The Manhattan plots depict the single nucleotide polymorphisms (SNPs) with the minor allele frequencies (MAF) above 0.05. The red line indicates the Bonferroni threshold of 7.55, which was determined by the –log_10_(*p*-value of 0.05/number of SNPs) that corresponds to -log_10_(0.05/1,753,576) = 7.55. **(C)** The genetic locus encompassing the SNPs associated with the QY max trait that was measured five days after salt treatment was examined in terms of its sequence divergence across the Arabidopsis accessions of the HapMap population. Open reading frames (ORFs) of the sequence that includes the SNPs associated with the QY max trait that was measured five days after salt treatment are depicted. This includes the missing data, insertions/deletions in the form of gaps, the sequence similarity across the sequenced HapMap accessions relative to Col-0 and the SNP positions (vertical dashed lines).

Furthermore, we attempted to develop a model for the salt tolerance index using rosette area while including the indices calculated across all 50 traits (**Figure S7**), but the model accuracy was very low. Compared to the multiple linear regression models (**Table S3**) and with an adjusted coefficient of determination (R^2^) values of 0.749 and 0.666 in control and salt stress conditions, respectively, the machine learning model resulted in a higher ranking of the photosynthetic parameters, which also had relatively high *p*-values in the multiple regression model (**Table S3**). Hence, the light-adapted chlorophyll fluorescence traits seem to be an important factor in plant development under control conditions, while the salt-induced decreases in the dark-adapted chlorophyll fluorescence traits might contribute to growth maintenance under salt stress.

### Characterising the genetic locus underlying maximum quantum yield

After performing the multivariate analyses, spatial correction was implemented for the GWAS on the raw phenotypic data according to a randomised split-plot block design (**Figure S2**). The significantly-associated SNPs from the single-trait GWAS models across all measured traits, time points and treatments have been summarised in **Tables S4** and **S5**. One notable association was found between the maximal quantum yield in the dark-adapted state (QY max) measured five days after salt treatment and SNPs on chromosome 5 (**Figure 5 A-B, Figure S8**). The association was specific to salt stress, as no significant associations were found using QY max that was measured under control conditions (**Figure 5 A**). The significant SNPs encompassed the promoter region of the constitutive photomorphogenesis protein 1 (COP1)-interacting protein 8 (CIP8 - AT5G64920) (Torii *et al*., 1999; Hardtke *et al*., 2002; Yuan *et al*., 2018) and the promoter regions of AT5G64930 and AT5G64940, which encode the regulator of pathogenesis related genes (CPR5) and an atypical protein kinase that is induced by heavy metal exposure (ABC1K8), respectively. A high level of genetic variation was observed across the HapMap accessions in this region when examining their divergence relative to Col-0 (**Figure 5 C**).

As no homozygous T-DNA insertion lines were identified for AT5G64930 and AT5G64940, we focused on exploring the role of CIP8 in the context of salt stress tolerance. To examine the role of CIP8 in maintaining QY max under salt stress, we used two homozygous T-DNA insertion lines (*at5g64920-1* and *at5g64920-2*, represented by SALK_138967 and SALK_023424, respectively, **Table S6**) and tested the CIP8 expression in both lines (**Figure S9, Table S6**). We then examined the salt-induced changes in rosette area and chlorophyll fluorescence (**Figure 6, Figure S10**). While both mutant lines developed larger rosettes under salt stress compared to Col-0 (**Figure 6 A-B**), this effect was specific to salt stress only in the case of *at5g64920-2*, and not *at5g64920-1*. Additionally, no differences were observed between the T-DNA insertion lines and Col-0 in terms of QY max (**Figure 6 C-D**) or in the light-adapted quantum yield (Fv′/Fm′) (**Figure 6 E-F**). Moreover, no significant differences were observed between Col-0 and T-DNA insertion lines in terms of non-photochemical (**Figure S10 A-B**) and photochemical quenching (NPQ and qP, respectively) (**Figure S10 C-D**). While these results suggest that CIP8 is involved in maintaining rosette area under salt stress, the role of CIP8 in regulating maximum quantum yield is still elusive. Additional genetic resources, such as overexpression lines, would be key in characterising this gene. In addition, developing homozygous T-DNA insertion lines in other candidate genes within the same locus (AT5G64930 and AT5G64940) might also contribute to understanding the maintenance of maximum quantum yield under salt stress, which was not feasible during this study.

**Figure 6.**
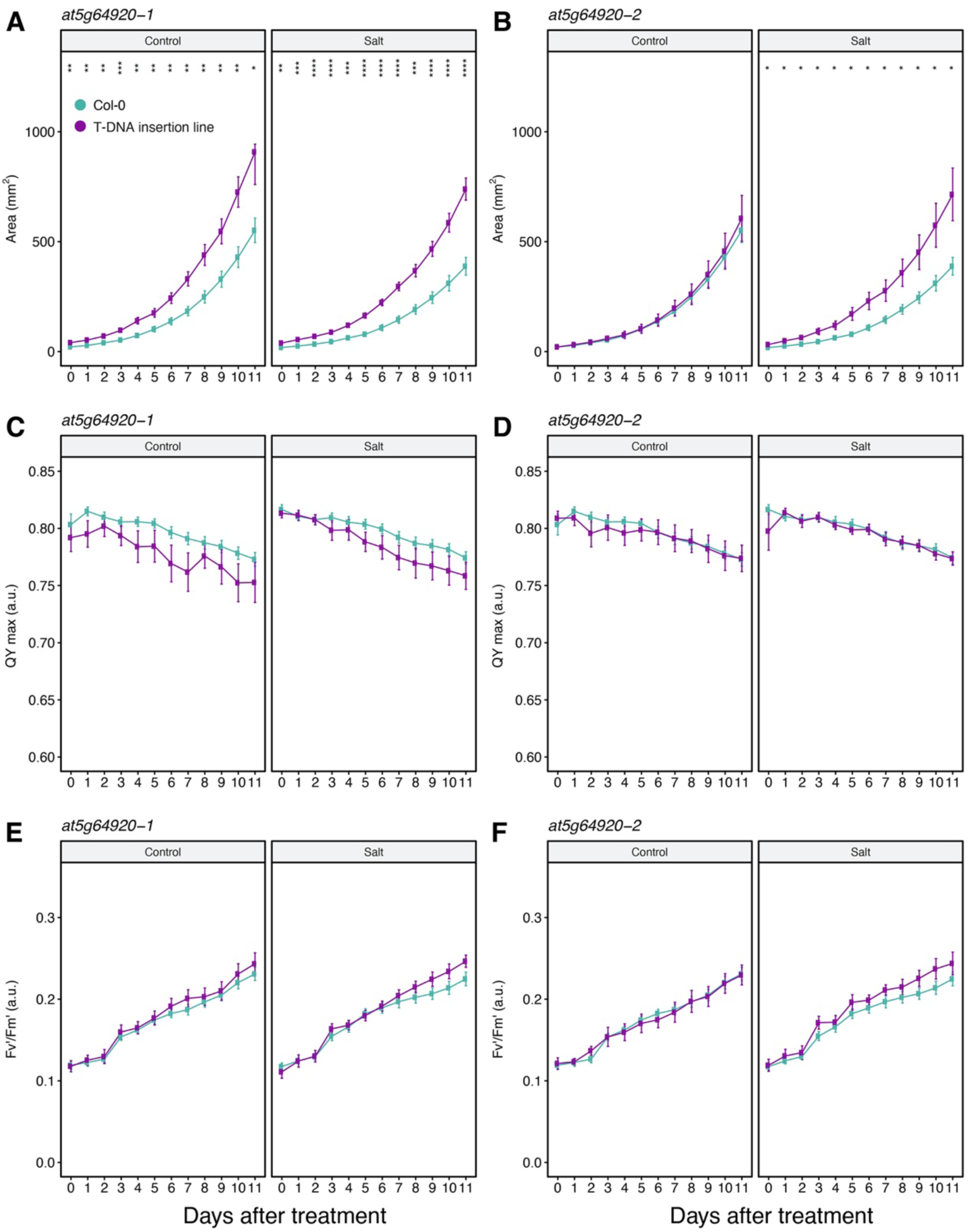
CIP8 mutants developed larger rosettes under salt stress. The gene AT5G64920 is annotated as the constitutive photomorphogenic 1 COP1-interacting protein 8 (CIP-8) and the salt stress responses of two mutant alleles of AT5G64920, i.e. *at5g64920-1* (SALK_138967) and *at5g64920-2* (SALK_023424) were examined for 11 consecutive days using high-throughput phenotyping. The two-week-old soil-grown plants were treated with a final concentration of 100 mM NaCl or control treatment, in which the change in the rosette area was monitored for 11 consecutive days. Comparisons of **(A,B)** rosette area, **(C,D)** maximum quantum yield (QY max) and **(E,F)** the light-adapted quantum yield (F_v_′/F_m_′) were conducted between the Col-0 background line and *at5g64920-1* **(A, C, E)** or *at5g64920-2* **(B, D, F)**, respectively. The asterisks above the graphs indicate significant differences between Col-0 and the T-DNA insertion mutant, where *, **, *** and **** indicate *p*-values < 0.05, 0.01, 0.001 and 0.0001, respectively, as determined by one-way ANOVA. The lines represent the mean calculated over 12 replicates per genotype and condition, while the error bars indicate the standard error of the mean.

**Figure 7.**
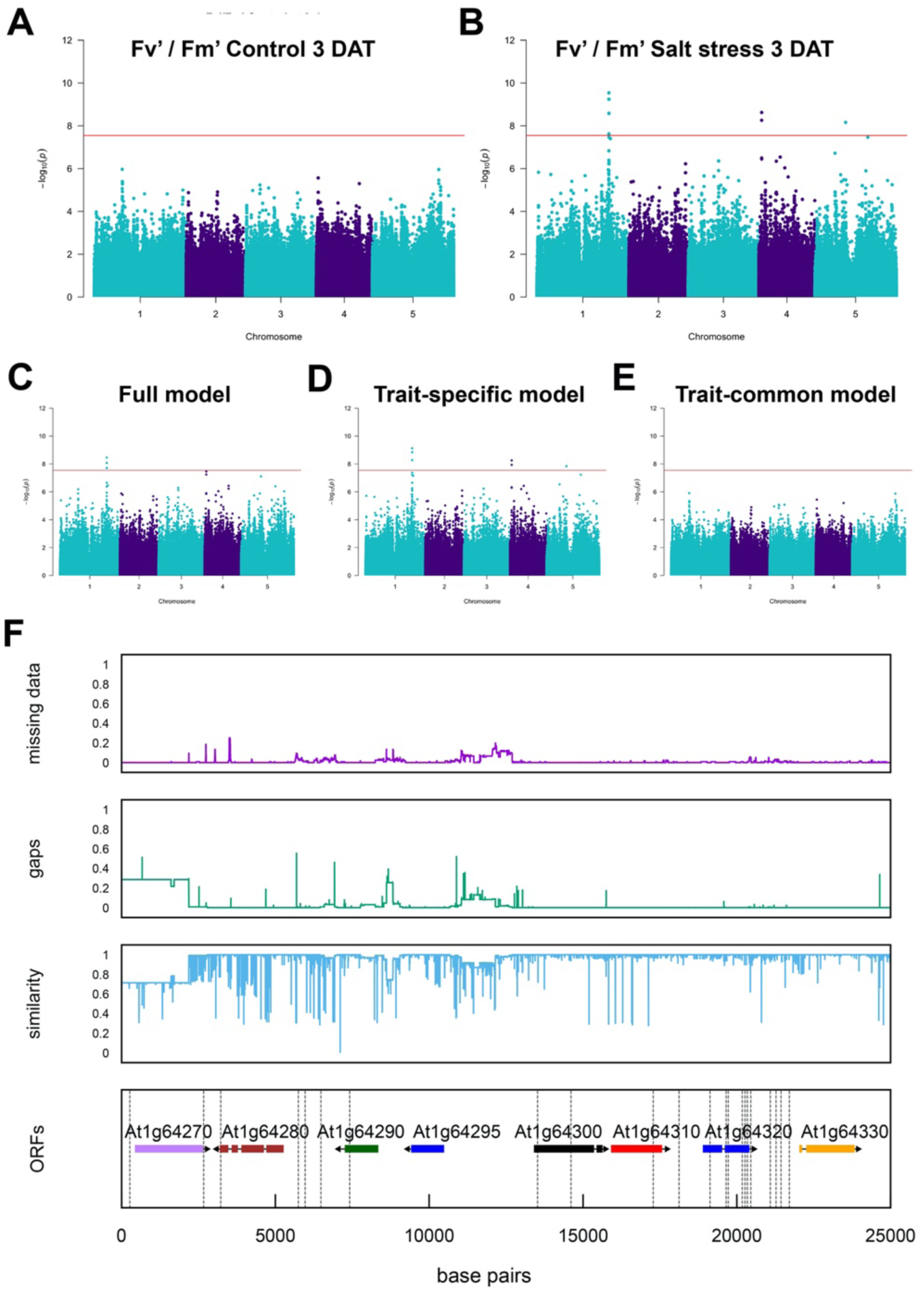
Association mapping of the light-adapted quantum yield. Single-trait genome-wide association study (GWAS) of the maximum quantum yield in the light-adapted state (F_v_′/F_m_′) using the highest photon irradiance of 440 μmol m^-2^ s^-1^ three days after treatment (DAT) under **(A)** control and **(B)** salt stress conditions. Multi-trait GWAS identified the genetic components contributing to the interaction between control and salt stress in F_v_′/F_m_′ that was measured three days after treatment. The Manhattan plots depict the single nucleotide polymorphisms (SNPs) with minor allele frequencies (MAF) above 0.05. The red line indicates the Bonferroni threshold of 7.55, which was determined by the –log_10_(*p*-value of 0.05/number of SNPs) that corresponds to -log_10_(0.05/1,753,576) = 7.55. The Manhattan plots also depict associations specific to **(C)** the interaction between genotype and the environment (MTMM full test), **(D)** environment (MTMM environment-specific test) and **(E)** genotype (MTMM environment - independent test), respectively. **(F)** Open reading frames (ORFs) of the sequence that includes the SNPs associated with the F_v_′/F_m_′ trait that was measured three days after salt treatment are depicted. This includes the missing data, insertions/deletions in the form of gaps, sequence similarity across the sequenced HapMap accessions relative to Col-0 and the SNP positions (vertical dashed lines).

### Light-adapted quantum yield under salt stress was compromised by SAPE

Single-trait GWAS using Fv′/Fm′ that was measured using the highest photon irradiance of Lss4 (440 μmol m^-2^ s^-1^) yielded several significant associations that were specific to salt stress (**Figure 7 A-B, Figure S11**). Chromosomes 4 and 5 contained a single significantly-associated SNP, while the 50 SNPs that were associated with Fv′/Fm′ on chromosome 1 between positions 23863584 and 23871063 were within linkage disequilibrium, therefore, were grouped into one locus. To examine whether these associations were truly specific to salt stress, an interaction GWAS model testing Fv′/ Fm′ that was measured on the same day under control versus salt stress was run using the MTMM GWAS (**Figure 7 C-E**). This model investigated the within-environment and between-environment variance by testing the genotypic (environment-independent), environmental (environment-specific) or genotype by environment (full test) effects on the association strength (Korte *et al*., 2012). The associations on chromosome 1, which were initially identified using single-environment GWAS, were again identified using the full and environment-specific models (**Figure 7 C-D**), but not in the environment-independent model (**Figure 7 E**). This suggested that the identified locus might play a role in regulating Fv′/Fm′ under salt stress and influenced the interaction between genotype and environment. However, the associations on chromosomes 4 and 5 were found only using the environment-specific models, which indicated that associations with SNPs on chromosome 1 were robust and salt-specific.

To further inspect the identified locus on chromosome 1, open reading frames and sequence divergence analyses were conducted to identify the hotspots of sequence diversity across the sequenced HapMap accessions (**Figure 7 F**). The majority of sequence diversity was localised between AT1G64280 and AT1G64295, but this region did not match the positions of the significantly-associated SNPs using Fv′/Fm′ (**Figure 7 F**), which were distributed along the promoter region of AT1G64270 and AT1G64280. AT1G64270 and AT1G64280 encode a transposable element and a regulator of the salicylic acid, respectively, while the coding region of AT1G64300 encodes an uncharacterised protein kinase. The highest concentration of SNPs that were significantly associated with Fv′/Fm′ was found between AT1G64310 and AT1G64320, which encode a pentatricopeptide repeat protein and an uncharacterised myosin heavy chain-like protein, respectively.

Subsequently, to test the involvement of each gene within the Fv′/Fm′ locus, we examined 12 homozygous T-DNA insertion lines (**Table S6**) and tested the insertion effect on gene expression (**Figure S12, Table S7**). All lines were examined for salt-induced changes in rosette size and chlorophyll fluorescence using the high-throughput phenotyping platform (**Figure 8, Figure S13**). Despite characterising the 12 homozygous T-DNA insertion lines in the coding sequence or promotor regions of the genes within Fv′/Fm′ locus, which can be found in the R-notebook (https://rpubs.com/mjulkowska/Arabidopsis_PSI_TDNA_pipeline_01), the only T-DNA insertion lines that showed a significant reduction in the target gene expression were *at1g64300-3* and *at1g64300-4* (**Figure S12**). These two mutant lines, *at1g64300-3* and *at1g64300-4*, developed larger rosettes under both control and salt stress conditions (**Figure 8 A-B**), with a more pronounced effect under salt stress, while the growth rate and growth maintenance had no significant changes (**Figure S14**). Nevertheless, both mutant lines showed an increase in the light-adapted quantum yield under salt stress compared to Col-0 (**Figure 8 C-D**), validating the GWAS association with this trait. Interestingly, both mutant lines also showed a reduced NPQ, specifically under salt stress, compared to Col-0 (**Figure 8 E-F**). However, no consistent differences were observed between the T-DNA insertion lines and Col-0 in terms of QY max or qP (**Figure S13**), indicating trait specificity of the locus association. Hence, AT1G64300 contributes to reduced plant size during salt stress, as salt-treated plants with reduced expression of this gene developed larger rosettes.

**Figure 8.**
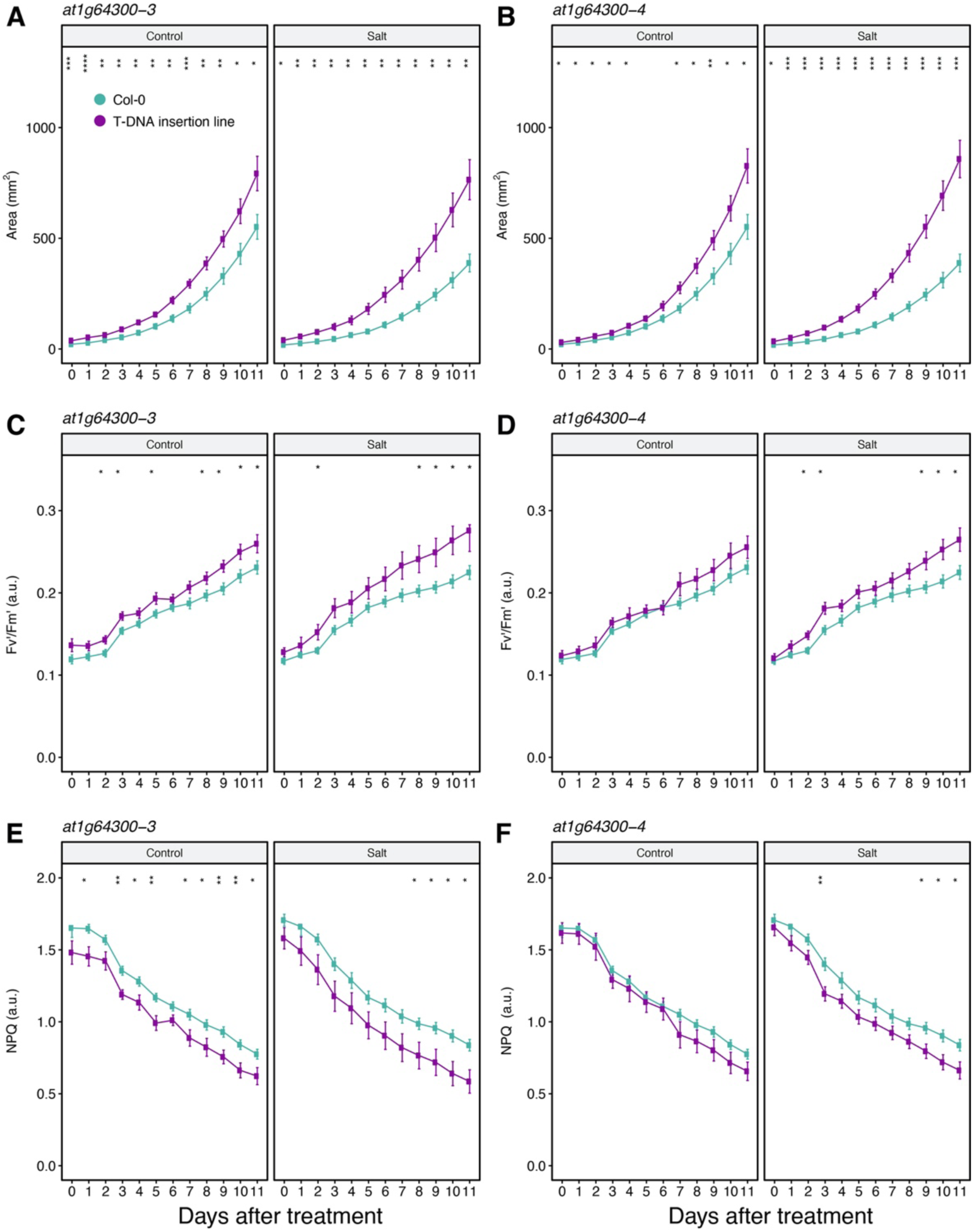
Mutants of the salt-affected photosynthetic efficiency (SAPE) gene in the F_v_′/F_m_′ locus demonstrate salinity tolerance. The salt stress responses of the two mutant alleles of AT1G64300, i.e. *at1g64300-3* (SALK_041569) and *at1g64300-4* (SALK_063928), were examined for 11 consecutive days using high-throughput phenotyping. The two-week-old soil-grown plants were treated with a final concentration of 100 mM NaCl or control treatment, in which the change in rosette size was monitored for 11 consecutive days. **(A,B)** Rosette area, **(C,D)** light-adapted quantum yield (F_v_′/F_m_′) and **(E,F)** non-photochemical quenching [NPQ = (F_m_ – F_m_′)/F_m_′)] values of the background line Col-0 (blue) and the mutants *at1g64300-3* and *at1g64300-4* (purple) are shown. The asterisks above the graphs indicate significant differences between Col-0 and the T-DNA insertion mutant, where *, **, *** and **** indicate *p*-values < 0.05, 0.01, 0.001 and 0.0001, respectively, as determined by one-way ANOVA. The lines represent the mean calculated over 12 replicates per genotype and condition, while the error bars indicate the standard error of the mean.

The AT1G64300 is an unknown protein kinase with two splice variants, in which the most dominant variant is 717 amino acids in length. Moreover, the predicted location of the serine/threonine kinase domain is towards the C-terminus (274 – 508), while the domain of the unknown function (DUF1221) is towards the N-terminus (21 – 237) (Willems et al., 2019). In addition, the AT1G64300 is predicted to contain up to 5 transmembrane domains in the positions 31-49, 108-128, 276-294, 329-350 and 435-453 (Hofmann & Stoffel, 1993), and it is predicted to be localised in the endomembrane systems, such as the Golgi apparatus or the endoplasmic reticulum (SUBAcon scores of 0.027 and 0.105), respectively, or in the plasma membrane (SUBAcon score of 0.815). Although the highest expression of At1G64300 was detected in the mature pollen (Schmid et al., 2005, Nakabayashi et al., 2005), it also expressed in the leaf and root tissue with a slight enhancement under multiple abiotic stresses, including heat, osmotic and wounding stress (Kilian et al., 2007). Due to the diverse nature of the promotor and coding sequence of AT1G64300, the representative haplotypes of this gene were not identified. Therefore, future studies focusing on the activity of this unknown kinase, its subcellular localisation and the generated phenotypes of the overexpression lines could provide insight into its function, which was outside of the scope of this study. Since the presence of AT1G64300 resulted in a reduced photosynthetic capacity under salt stress, we opted to name AT1G64300 as the salt-affected photosynthetic efficiency (SAPE) gene.

## Discussion

Salt stress effects have been primarily explored in terms of the ionic component, where increased ion accumulation in the shoot can reduce biomass accumulation and lead to premature senescence. However, growth rate reductions typically occur before significant sodium ions accumulation in the shoot (Fricke *et al*., 2006, Rajendran *et al*., 2009; Adem *et al*., 2014; Tilbrook *et al*., 2017). In addition, ion exclusion does not reflect salt stress tolerance across diverse plant species (Genc *et al*., 2007). More recently, salinity studies have begun investigating new traits that can potentially contribute to salinity tolerance (Negrao *et al*., 2017, Morton *et al*., 2019, van Zelm *et al*., 2020), including stress signaling pathways (Choi *et al*., 2014; Evans *et al*., 2016), cell wall remodeling (Feng *et al*., 2018), transpiration use efficiency (Al-Tamimi *et al*., 2016) and root system architecture (Julkowska *et al*., 2017), thus, breaking down salinity tolerance into more genetically tractable components (Morton *et al*., 2019). The non-destructive methods developed in high-throughput phenotyping allowed the recording of more sophisticated traits (Awlia *et al*., 2016), providing simultaneous and multifaceted understanding of plant size, architecture and photosynthetic efficiency during plant development and in response to environmental cues (Negrão and Julkowska, 2020). In this study, we used high-throughput phenotyping to enable in-depth GWAS and have identified over 1,000 unique SNPs that were associated with the responses of Arabidopsis to salt stress (**Table S3, Table S4**). The GWAS associations identified for a single trait and time point were strengthened by the time-series approach, where a repeatedly significant association across time and traits provided higher confidence in the association. This layered association approach highlights the genetic constituents of the processes that occur temporally, which could be overlooked in single-time point GWAS. Although we did not identify a single locus that was common across all traits or time points, the identified associations in this study are a valuable resource for investigating plant responses to salt stress as well as plant development under control conditions.

In this work, we observed that salt stress caused rapid changes in rosette area, colour and the chlorophyll fluorescence traits of Fv′/Fm′, NPQ and qP (**Figure 1, Figure S3 - S4**), which were also observed in nine diverse Arabidopsis accessions (Awlia *et al*., 2016), suggesting that Fv′/Fm′, NPQ and qP could be reliable markers for early salt stress responses. Rosette morphology, QY max and Ft, on the other hand, were affected at a later stage after salt treatment (**Figure 1, Figure S3**). This distinction between changes in traits that reflect early and late responses to salt stress supports the hypothesis of two distinct phases of osmotic and ionic responses (Munns and Tester, 2008). In addition, we observed a wide range of rapid salt stress responses, including changes in rosette colouring and photosynthetic capacity, resulting in the rapid reduction of light-adapted quantum yield (**Figure 1 D**). Moreover, we observed a more severe inhibition of growth in the second interval (**Figure 2 A**), corresponding to the ionic phase of salt stress (Munns and Tester, 2008). Interestingly, the correlations between the salt tolerance indices across the two intervals (**Figure 2 B**) were found to be remarkably low. This suggests that the maintenance of growth could be regulated by different genetic components during the two phases of salt stress exposure.

No robust genetic signals associated with growth rates or salt tolerance indices were detected in our GWAS (**Figure S6**), despite salt tolerance indices highlighting potential yield maintenance loci in barley under salt stress, which overlapped with yield and harvest index quantitative trait loci (QTL) regions (Saade *et al*., 2016). In the barley study, the data was collected at one time point and across two field trials, which helped strengthen the statistical power and improved the detection of genetic signals. However, the salt tolerance indices presented here (**Figure 2**) were estimated using growth rates that were calculated across three-day intervals and derived from a single experiment. Similarly, mapping the relative growth rate under salt stress across four designated intervals using a rice diversity panel showed little success in detecting QTLs, in contrast to transpiration use efficiency, which was associated with multiple QTLs (Al-Tamimi *et al*., 2016). Furthermore, several studies have successfully identified QTLs underlying the natural variation in the root growth rate (Meijón *et al*., 2014; Slovak *et al*., 2014). However, these studies were performed at a much smaller timescale with only one condition. While salt tolerance indices can be useful to evaluate the relative performance of different genotypes (Morton *et al*., 2019), our results suggest that the shoot growth rate and the salt tolerance indices are genetically complex traits and may require a high number of replicates and a larger mapping population for effective GWAS.

The power of high-throughput phenotyping lies in measuring a large number of traits through time, including traits not suspected to contribute to stress tolerance, which provides an unbiased approach. This data richness provided us with the opportunity to not only identify new genetic components of salinity tolerance, but also to improve our understanding of the relationships between different traits. For this purpose, we deployed the correlation analysis (**Figure 2**) and the machine learning models (**Figure 3**) to identify which traits have the highest contribution to plant size under control and salt stress conditions. Although machine learning and statistical data-mining have been recognised to reduce the complexity of high-throughput phenotyping experiments (Rahaman *et al*., 2019), there are only a handful of examples that used these methods besides multispectral imaging and image processing (Lee *et al*., 2018; Gao *et al*., 2019; Gutiérrez *et al*., 2016; Ubbens and Stavness, 2017; Grinberg *et al*., 2020). Both the correlation analysis and the gradient-boosted tree models have highlighted QY max as an important contributor to rosette area under control and salt stress conditions (**Figure 2**, **Figure 3**). As QY max is known to be a robust parameter of photosynthesis, it is only compromised with prolonged exposure to stress (**Figure 1**)

(Kalaji *et al*., 2011), which reflects the photoinhibition and inactivation of the photosystem II reaction center (Murata *et al*., 2007). In our study, only a single locus underlying QY max was identified under salt stress (**Figure 5**) and the examined T-DNA insertion lines within this locus did not differ from Col-0 in terms of QY max (**Figure 6, Figure S10**). As only a small number of accessions showed compromised QY max under salt stress (**Figure 1 C**), this could have compromised the statistical power of the GWAS model, limiting the detection of other genetic constituents contributing to the maintenance of QY max. The gradient-boosted trees model identified several light-adapted chlorophyll fluorescence parameters as important contributors to rosette area, specifically under salt stress (**Figure 3**). The importance of these traits would have been overlooked in the multiple linear regression models (**Table S3**). Hence, the importance of the light-adapted chlorophyll fluorescence in the machine learning model was in line with the observation that these traits change at earlier time points and are more pronounced when compared to the dark-adapted chlorophyll fluorescence traits. The contributions of each of these traits to overall salinity tolerance would require more detailed analyses of the underlying genetic players that we have identified here (**Table S4**), however, this in-depth analysis is beyond the scope of this work.

As Fv′/Fm′ showed early changes in response to salt stress, especially when combined with multiple light-adapted chlorophyll fluorescence traits, we opted to examine its genetic constituents in more detail (**Figure 7**). In addition, the MTMM GWAS approach allowed us to test the interaction between Fv′/Fm′ measured under control and salt stress (Korte *et al*., 2012), highlighting the locus on chromosome 1 as a salt-specific association. Next, the detailed study of the T-DNA insertion mutants of genes located within the Fv′/Fm′ locus (**Figure 8, Figure S13**) showed that the reduced expression of AT1G64300, which encodes a protein kinase of unknown function, resulted in the improved maintenance of Fv′/Fm′ under salt stress. The other candidate genes within the Fv′/Fm′ locus, AT1G64310 and AT1G64320, that encode a pentatricopeptide repeat protein and a myosin heavy chain-like protein, respectively, may also contribute to the maintenance of Fv′/Fm′ under salt stress. However, no homozygous T-DNA insertion lines that displayed a reduction in the expression of AT1G64310 and AT1G64320 were identified (**Figure S12**). Interestingly, the T-DNA insertion lines of AT1G64300 developed larger rosettes and maintained higher Fv′/Fm′ under salt stress than Col-0, suggesting that the functional protein kinase could be hindering growth maintenance in saline conditions. Therefore, we named this protein kinase as the salt-affected photosynthetic efficiency (SAPE) gene. Although SAPE’s function remains unknown, the available transcriptome data reveals that its expression is slightly upregulated in response to multiple abiotic stresses, including heat, genotoxic and osmotic stress (Kilian *et al*., 2007). Moreover, the closest SAPE homologue in the Arabidopsis genome is AT5G41730 that encodes an unknown protein kinase, but has a low expression similarity (Waese *et al*., 2017), making it unlikely that the two genes have redundant functions. Single T-DNA insertion mutants, with reduced SAPE expression, have indeed shown detectable differences relative to the Col-0 background line in terms of their Fv′/Fm′ and rosette area under salt stress, suggesting that SAPE might not have homologues with redundant functions. Furthermore, the specificity of the SAPE phenotype is also noteworthy, as both mutant lines differed from Col-0 in terms of their Fv′/Fm′ and NPQ values (**Figure 8**), which are closely linked to quantum yield, but not in their QY max or qP values (**Figure S13**). Hence, this indicated that SAPE can affect the responses to salt stress by acting exclusively on the light-adapted chlorophyll fluorescence traits. As dissecting the molecular mechanisms of SAPE was beyond the scope of this work, future studies of SAPE will help to better understand the maintenance of photosynthetic efficiency under salt stress.

To summarise, our work presents an extensive overview of the salt stress responses of 191 Arabidopsis accessions. In this study, we were able to classify the different traits as early and late salt stress responses, which can correspond to the osmotic and ionic phases of salt stress (*sensu* Munns & Tester, 2008). Furthermore, the identified Fv′/Fm′ locus seems to reflect the early changes in chlorophyll fluorescence, where the SAPE mutants better maintained their quantum yield and rosette size under salt stress relative to the wild-type and control-treated plants throughout the experiment, suggesting that targeting the genetic constituents of the early responses to salt stress could improve salinity tolerance. Hence, further investigations into the wealth of candidate genes that were highlighted in this study can act as useful markers for breeding salt tolerant crops in arid and semi-arid regions to support global food security.

## Acknowledgements

We thank King Abdullah University of Science and Technology (KAUST) for funding this work, and Photon Systems Instruments (PSI) for their scientific and technical support. Special thanks to Dr. Xavier Sirault (CSIRO, Australia) for suggesting the use of the saturation method for salt imposition, Radka Mezulániková and Jaromír Pytela (PSI, Czech Republic) for assisting with the phenotyping and Prof. Christa Testerink (UVA, Netherlands) for providing the HapMap seeds. We also thank the KAUST Greenhouse Core-lab personnel: Dr. Richard Soppe, Mr. Mupalla Reddy, Mr. Gomerito Sagun, Mr. John Rahmer, Mr. Thomas Hoover and Mr. Johnard Balangue for their help in propagating the plant material that was used in this study and for running the high-throughput phenotyping facility at KAUST.

---

**Table S1. The examined Arabidopsis accessions.** Details of the accession Nordborg ID, the seed collectors’ names and the latitude and longitude coordinates of the seed collection sites were obtained from the 1001 Arabidopsis Genomes Project, while the accession CS numbers were obtained from the Arabidopsis Information Resource (TAIR) website.

**Table S2. Summary of the [red:green:blue] (RGB), greenness and chlorophyll fluorescence trait descriptions, values and significant changes.** The plants were screened by the high-throughput phenotyping system from one hour to seven days after control or salt treatment. Mean values were based on the last day of measurement of three or more biological replicates per genotype and condition using standard error. Greenness hues were represented as [red:green:blue] coordinates in the RGB space, which were obtained from the RGB image colour segmentation process. The salt-induced significant changes and the days at which the changes were observed relative to control conditions were determined using one-way analysis of variance for all days after transfer (DAT).

**Table S3. Multiple linear regression models for rosette area.** All the measured traits were implemeted as explanatory variables, except for rosette perimeter and the greenness hues, in the multiple linear regression models of the rosette area (mm^2^). The term ‘Estimate’ represents the model slope values ascribed to each trait, while the standard error (SE) reflects the coefficient estimate’s variation from the response variable’s actual value. The t-value represents the coefficient estimate’s deviations from 0, while Pr(>|t|) is the probability of observing any value equal or larger than t. The significance of the values is included, where ‘ ’, ‘.’, *, ** and *** indicate *p*-values > 0.1, < 0.1, 0.05, 0.01, and 0.001 respectively.

**Table S4. Genome-wide association study (GWAS) results of the significantly-associated loci.** The associations were performed using the spatially-corrected Arabidopsis phenotypic data derived from the RGB and chlorophyll fluorescence images consisting of rosette size, morphology, greenness and chlorophyll fluorescence traits that were captured under control and salt stress conditions. The salt tolerance index (STI) was estimated in the two time intervals of 0 to 3 and 4 to 7 days after treatment. Single-trait and multi-trait mixed models (MTMM) were used in the GWAS, in which the logarithm of odds (LOD) score was estimated by -log_10_(*p*-value). The explained variance refers to the natural variation observed in the studied population.

**Table S5. Summary of the identified loci that were generated from the genome-wide association study (GWAS).** The significantly-associated SNPs across multiple time points and traits were compiled, while highlighting the salt-specific traits. The single-trait GWAS model was used to identify all the listed associations, where each significant SNP that is located within 10 kbp of another significant SNP was considered in linkage disequilibrium and treated as one locus. Upstream and downstream SNPs refer to the flanking SNPs of each estimated locus. The logarithm of odds (LOD) score was estimated by -log_10_(*p*-value), while the number of SNPs, traits, days and treatments of each locus has been indicated. Treatment number values of 1 indicate associations found in either control or salt treatment, 2 indicates both treatments, while 3 indicates associations common across the two treatments and using the ratio of salt over control. The matrix illustrates the significant associations in each locus per trait using a binary code of 0 and 1, reflecting the absence or presence of a significant association, respectively. The traits that were included in the matrix have been labeled as treatment_trait_day.

**Table S6. The T-DNA insertion lines and primers that were used for the genotypic validation.** The primers were designed by the T-DNA Express: Arabidopsis Gene Mapping Tool of the Salk Institute Genomic Analysis Laboratory, where Col-0 (wildtype) was used as the background line for generating the T-DNA insertion lines. The left and right primers (LP and RP, respectively) used for validating the T-DNA insertion lines have been listed as well as the expected band size of Col-0 (WT), which was obtained through the PCR reaction of LP and RP. The expected size of the band in the T-DNA insertion mutant, which was obtained through the PCR reaction of RP and the SALK or SAIL-specific border primer (BP) has also been listed.

**Table S7. The primers that were used for the target gene expressions of the T-DNA insertion lines.** Fwd and Rev refer to the forward and reverse primer sequence directions, respectively.

**Figure S1.**
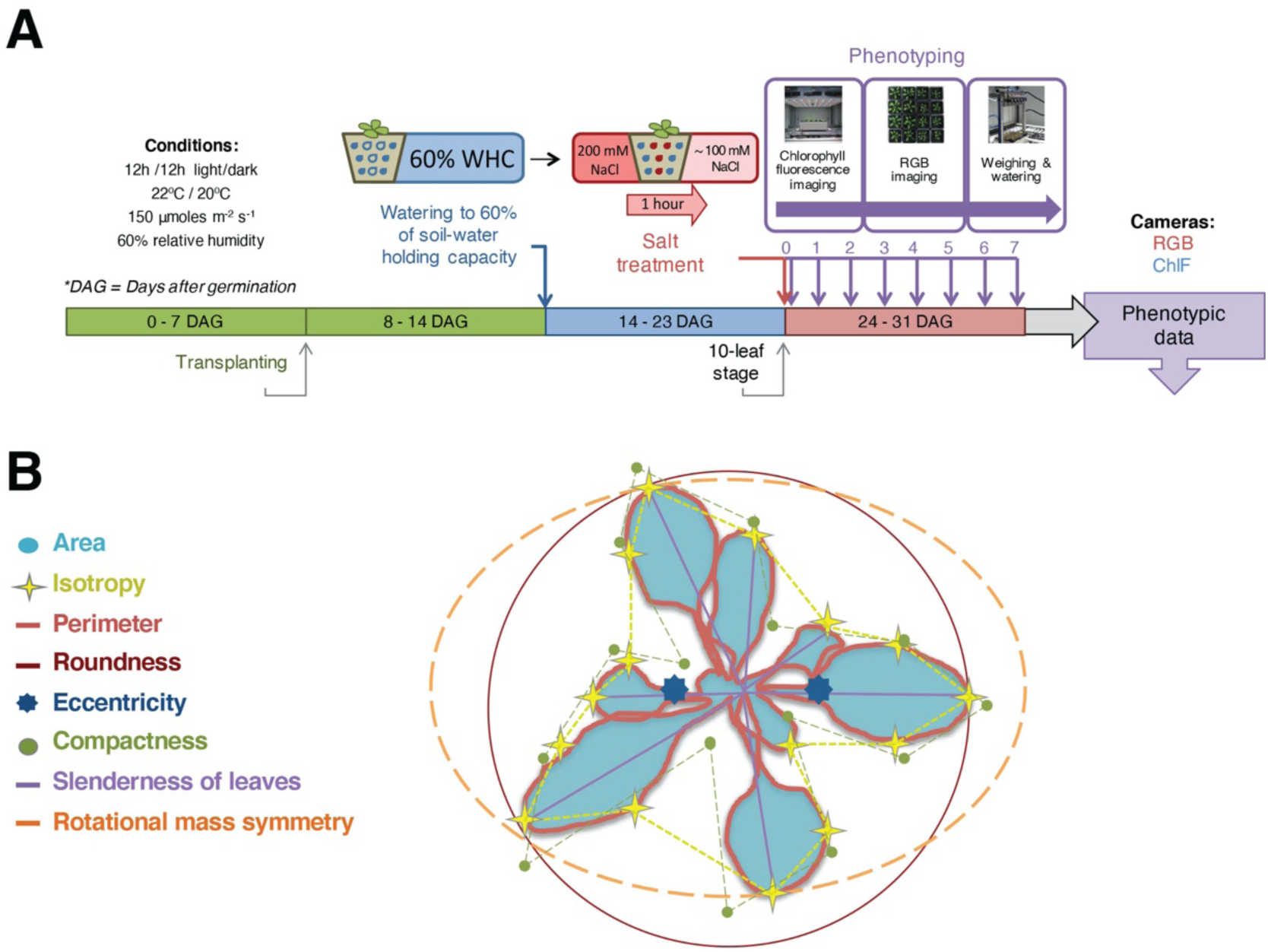
Illustrations of the salt treatment, phenotyping protocol and the traits used for monitoring the early responses to salt stress. **(A)** Arabidopsis accessions were germinated in a growth chamber at a 12h/12h, 22°C/20°C, light/dark regime with 60% humidity. Seedlings of similar size were transplanted into freshly sieved soil seven days after germination (DAG). At 14 DAG, watering was controlled to reach 60% of the soil-water holding (SWH) capacity. Salt treatment was applied to the seedlings at the 10-leaf stage (24 DAG) by saturating the pots for one hour in 250 mM NaCl, resulting in an estimated final concentration of 100 mM NaCl. Control and salt-treated plants were phenotyped after one hour and through seven consecutive days using the high-throughput PlantScreen^TM^ conveyor system (PSI, Czech Republic). Chlorophyll fluorescence (ChlF) imaging, [red:green:blue] (RGB) imaging and automated weighing were performed as part of the phenotyping setup. **(B)** Schematic overview of the parameters captured using the RGB imaging reflecting rosette size and morphology.

**Figure S2.**
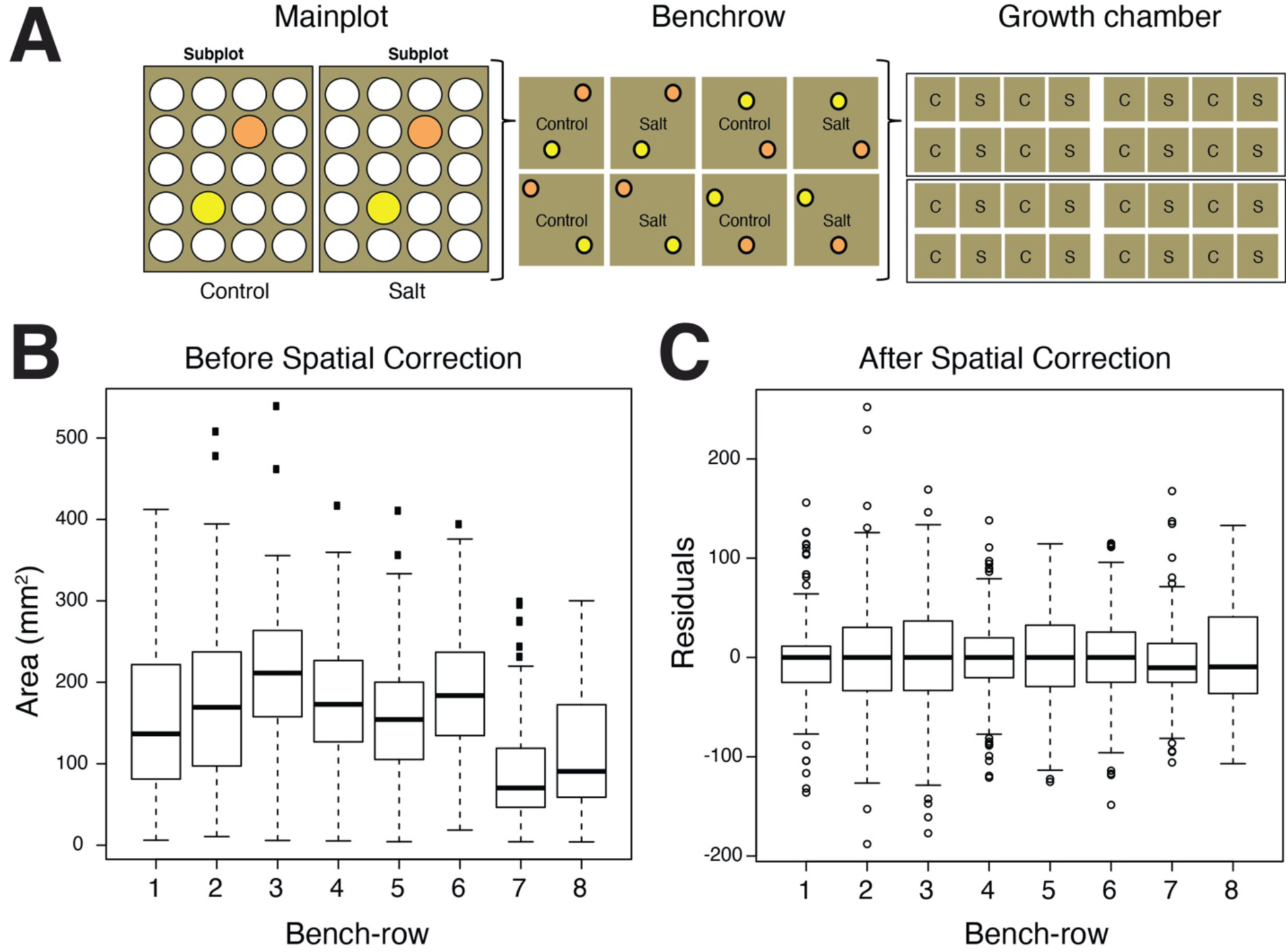
Experimental design and spatial correction rationale. **(A)** The split-plot block design accounted for the pot position (sub-plot), the position of the control and salt-treated trays (main-plot), the four pairs of trays on a bench (bench-row) and the position within the growth chamber. Spatial correction was performed using the ASreml package in R to account for the experimental design and possible spatial effects of position, microenvironments and shading on the recorded phenotypic data. Total rosette area (mm^2^) per bench-row (n=8) measured on the final day of phenotyping (seven days after treatment) **(B)** before and **(C)** after spatial correction.

**Figure S3.**
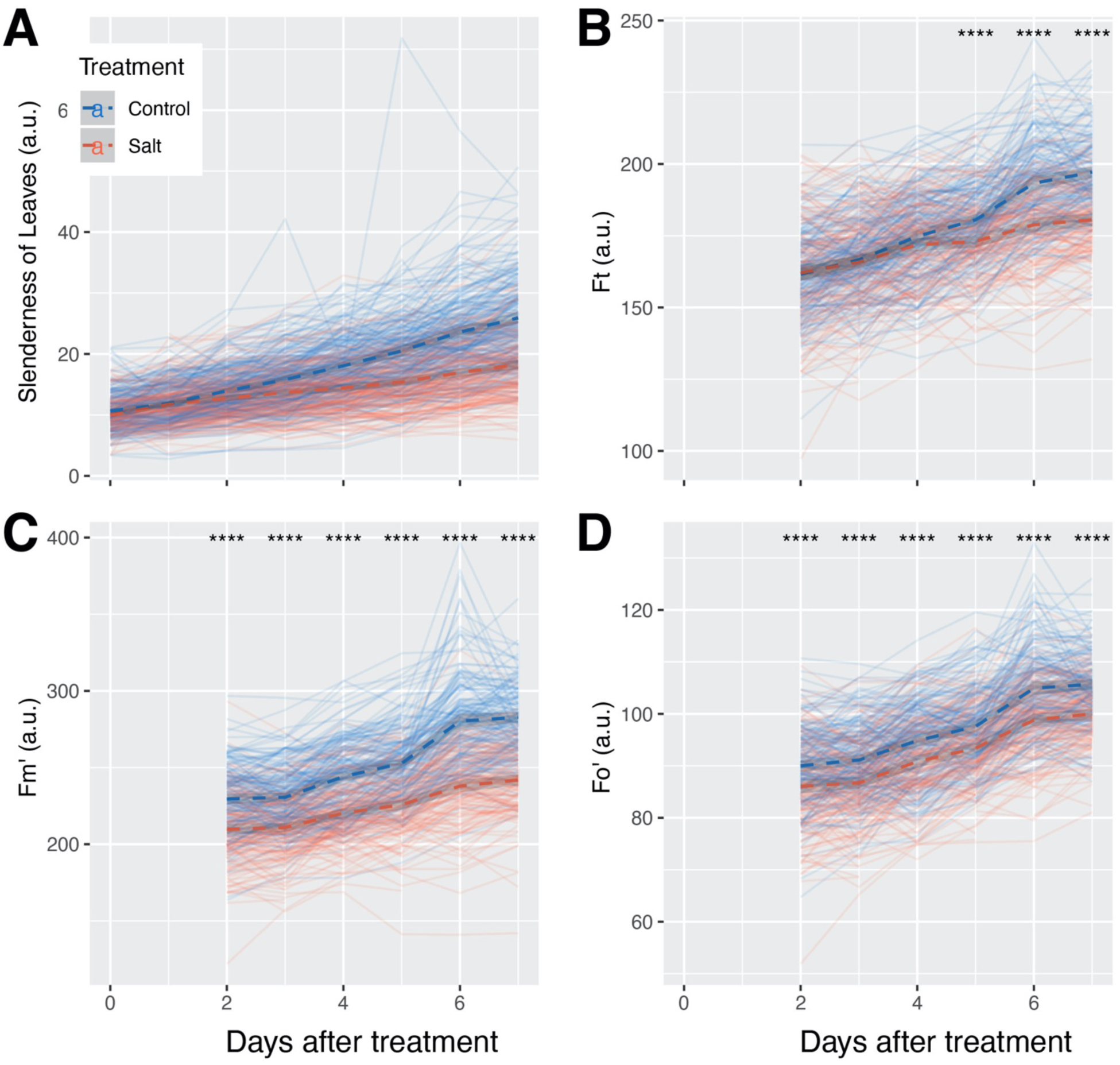
Additional plant morphology and photosynthetic efficiency traits altered by salt stress. The salt stress responses of three-week-old soil-grown plants that were treated with a final concentration of 100 mM NaCl were examined as described by (Awlia *et al*., 2016). **(A)** Slenderness of leaves, which is the ratio between area and distance from the plant center to its leaf tips, was calculated based on the pixels captured by the [red:green:blue] (RGB) camera immediately after salt treatment and seven days thereafter. **(B)** Instantaneous fluorescence (Ft), **(C)** maximum chlorophyll fluorescence (Fm′) and **(D)** minimum chlorophyll fluorescence (Fo′) in the light-adapted state were only captured two days after salt treatment (due to technical difficulties) using the highest photon irradiance (440 μmol m^-2^ s^-1^) with the light curve protocol of chlorophyll fluorescence imaging. Each line represents the trajectory per accession through time, where blue and red lines indicate plants grown under control or salt stress conditions, respectively. The dashed lines represent the mean of the population per condition and trait, while the grey band represents the confidence interval. The asterisks above the graphs indicate significant differences between control and salt-treated plants within the entire population, where *, **, *** and **** indicate *p*-values < 0.05, 0.01, 0.001 and 0.0001, respectively, as determined by one-way ANOVA.

**Figure S4.**
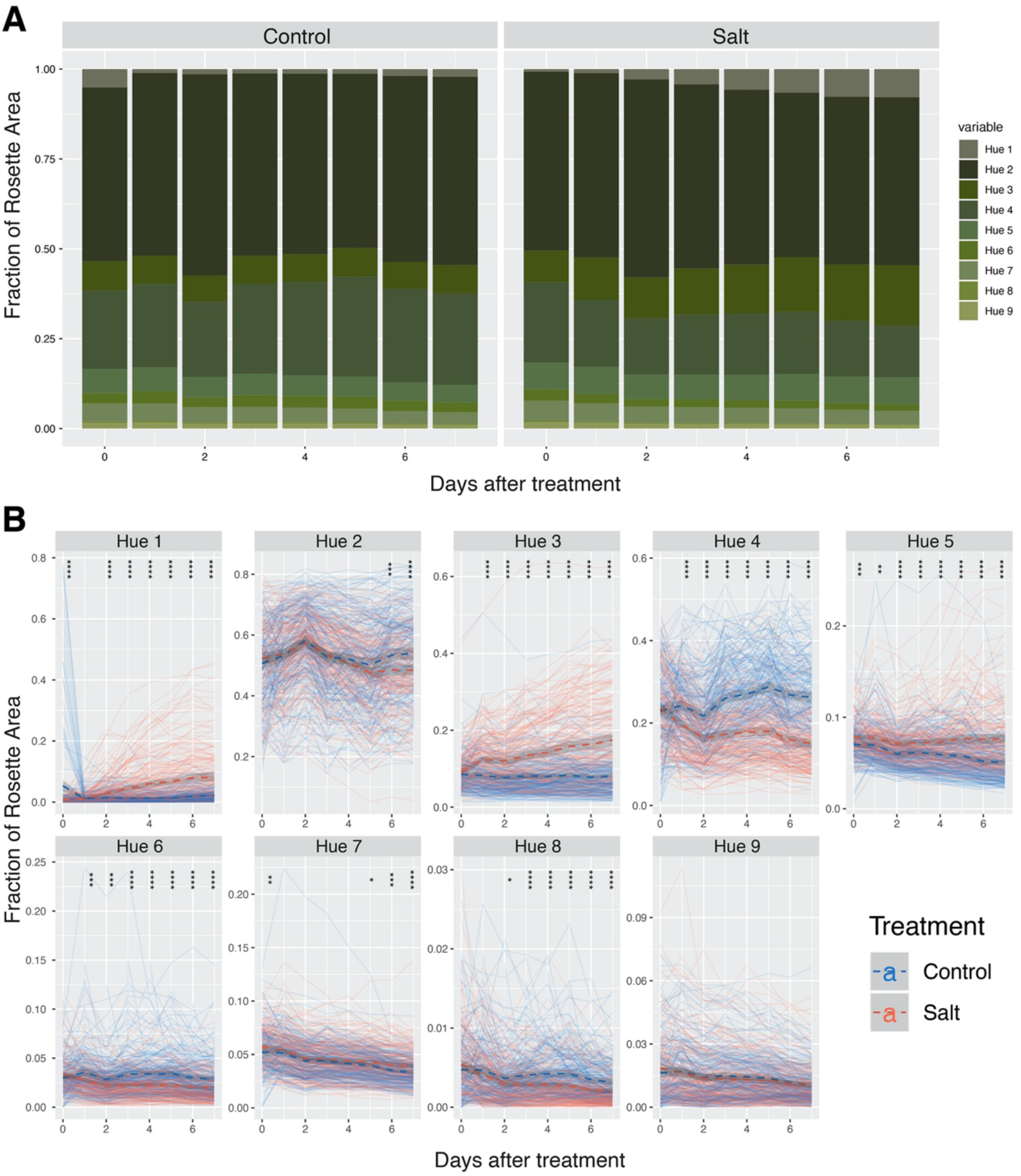
Changes in rosette greenness were affected by both plant development and salt treatment. The salt stress responses of three-week-old soil-grown plants treated with a final concentration of 100 mM NaCl were examined as described by (Awlia *et al*., 2016). The colour pixels of the Arabidopsis rosette [red:green:blue] (RGB) images were clustered into the nine most abundant green hues based on k-means clustering. **(A)** The relative abundance of each greenness hue is shown as a fraction of rosette area that was measured through time under control (left) and salt stress (right) conditions, where the values represent the grand mean of the population. **(B)** Comparisons of the relative hue abundance among the Arabidopsis accessions grown under control and salt stress conditions are shown. Each line represents the relative hue abundance per accession through time, where blue and red lines indicates plants grown under control and salt stress conditions, respectively. The dashed lines represent the mean of the population per condition and trait, while the grey band represents the confidence interval. The asterisks above the graphs indicate significant differences between control and salt-treated plants within the population, where *, **, *** and **** indicate *p*-values < 0.05, 0.01, 0.001 and 0.0001, respectively, as determined by one-way analysis of variance. Note that the hues were plotted using different scales to aid the readability of the graphs.

**Figure S5.**
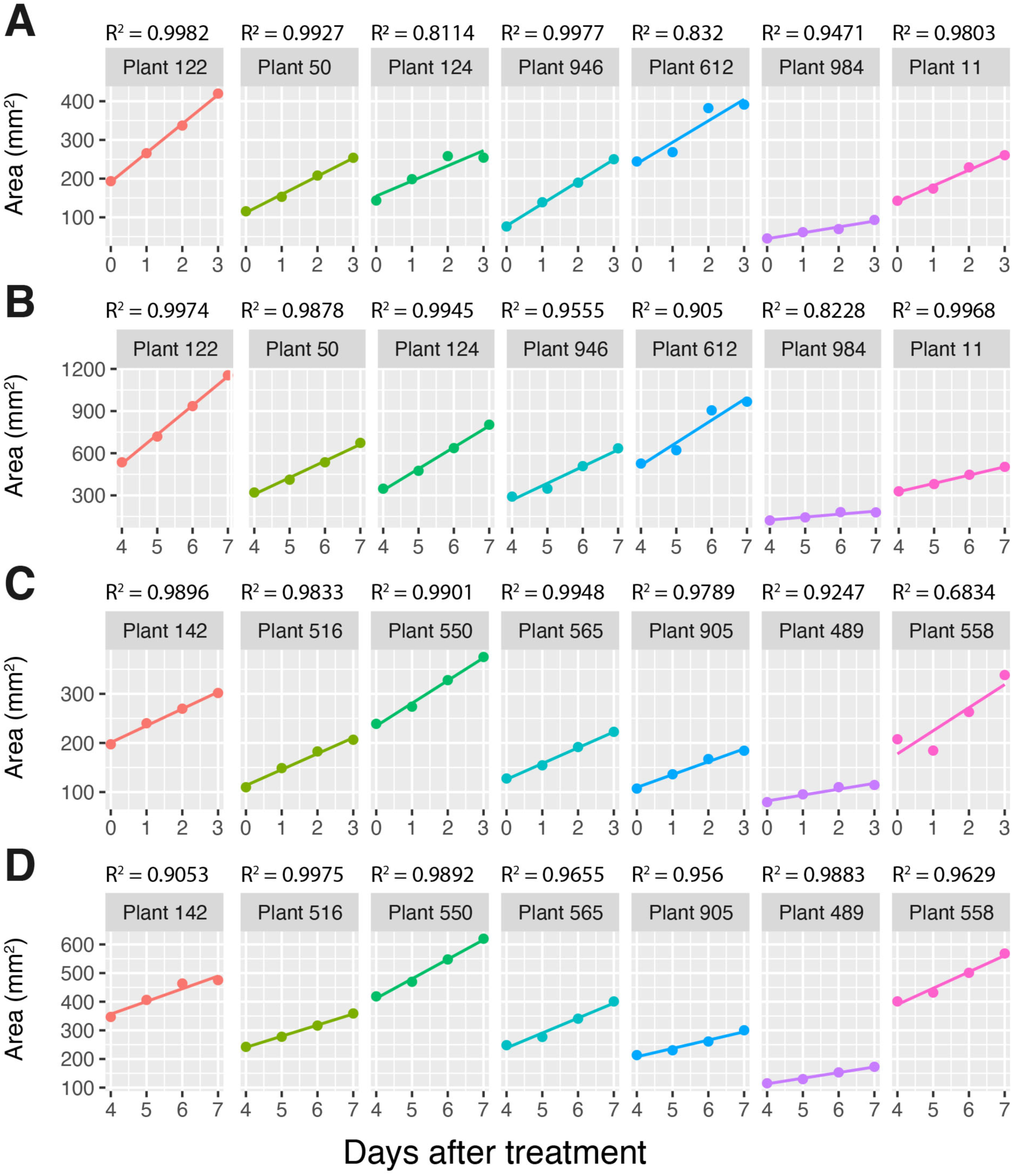
Fitting a linear function to the change in rosette area over two time intervals. The growth rate per plant was estimated by fitting a linear function using the lm() command in R to the change in rosette area (mm^2^) over intervals 1 and 2 (0 to 3 and 4 to 7 days after treatment, respectively) under control **(A-B)** and salt stress **(C-D)** conditions. The scatter plots indicate the change in rosette area in seven randomly selected (A-B) control or (C-D) salt-treated plants. The solid lines indicate the linear model (lm), while the adjusted coefficient of determination (R^2^) values that reflect the model fitting, can be found above each graph.

**Figure S6.**
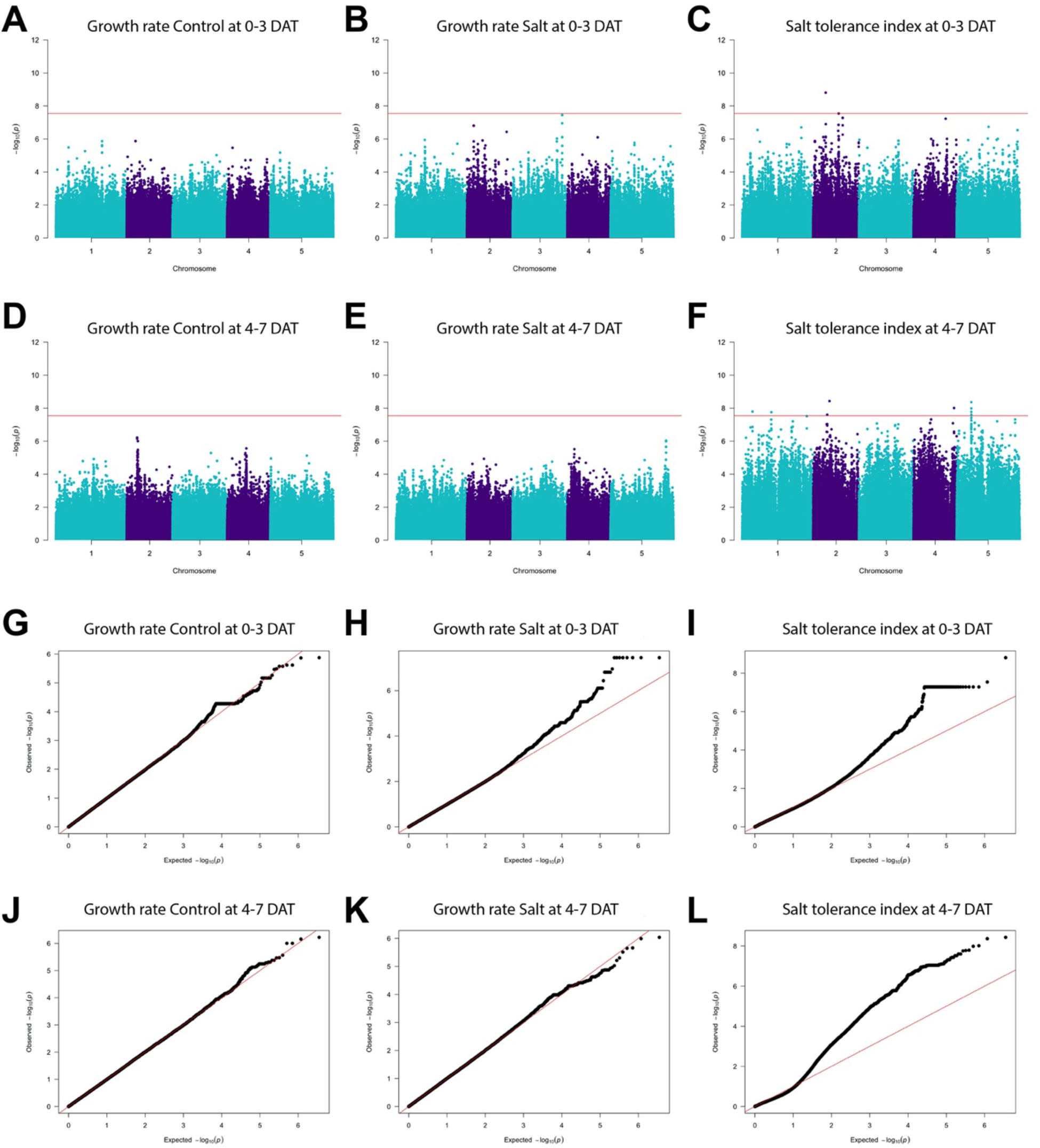
Identifying the genetic components underlying growth during intervals 1 and 2, reflecting the early and late responses to salt stress. Single-trait genome-wide association studies (GWAS) was performed on the growth rate derived from the change in rosette area from 0 to 3 and 4 to 7 days after treatment (DAT) under **(A,D)** control and **(B,E)** salt stress conditions as well as on **(C,F)** the salt tolerance index (STI). STI was calculated by dividing the growth rate of each accession measured under salt stress by the growth rate measured under control conditions. The Manhattan plots depict the single nucleotide polymorphisms (SNPs) with minor allele frequencies (MAF) above 0.05. The red line indicates the Bonferroni threshold of 7.55, which was determined by the –log_10_(*p*-value of 0.05/number of SNPs) that corresponds to -log_10_(0.05/1,753,576) = 7.55. Quantile-quantile (Q-Q) plots were generated to reflect the single-trait GWAS fitting of the growth rates in interval 1 (0 to 3 DAT) under **(G)** control, **(H)** salt stress and **(I)** STI-interval 1 as well as in interval 2 (4 to 7 DAT) under **(J)** control, **(K)** salt stress and **(L)** STI-interval 2.

**Figure S7.**
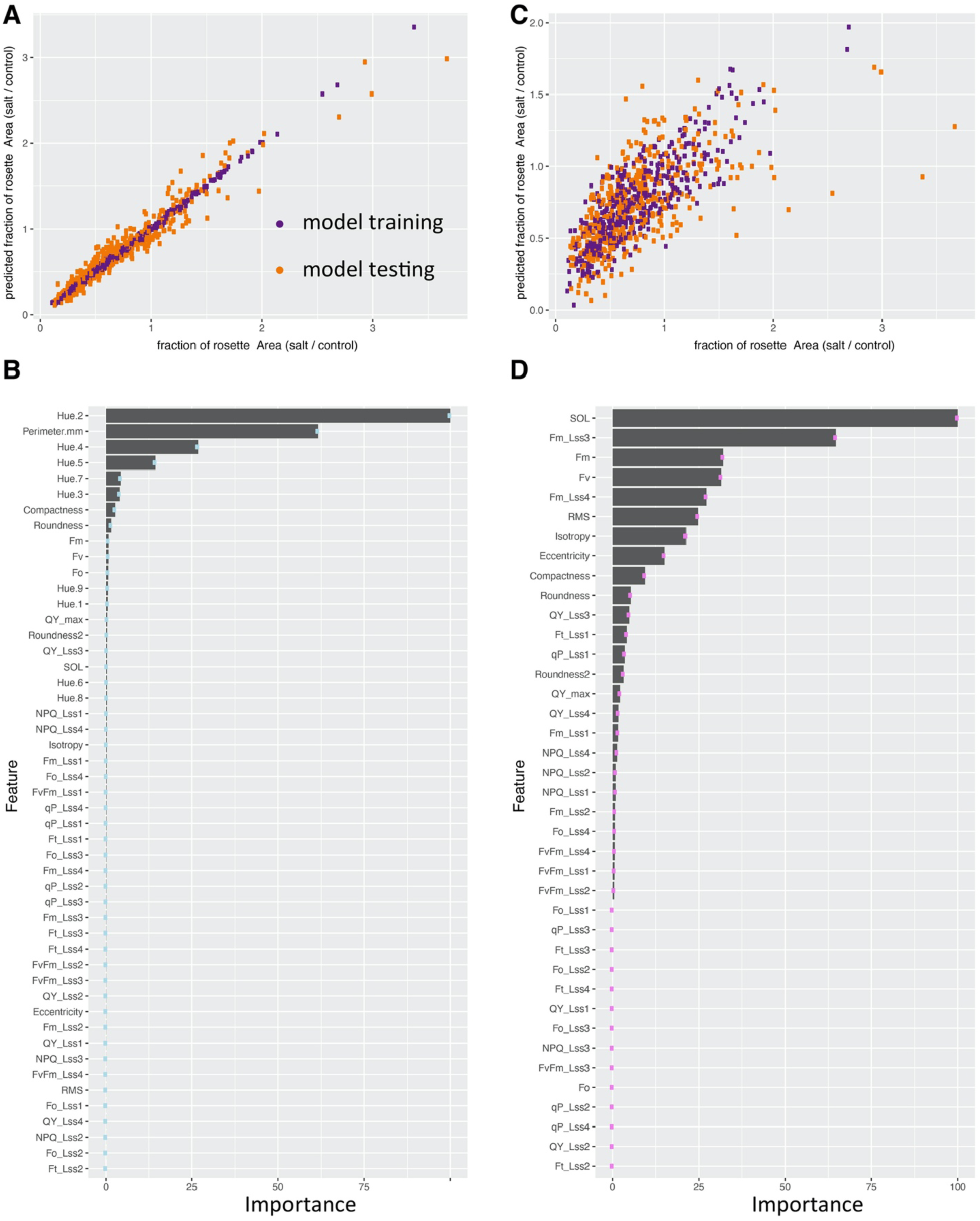
Machine learning using the salt tolerance index can identify the traits that contribute the most to the salt-induced changes in rosette area. **(A)** The gradient-boosted tree model was developed using 50% of the phenotypic data to predict the salt-induced changes in rosette area. This was done by applying the salt tolerance index (STI) formula (area_salt_/area_control_) with respect to all the measured traits (trait_salt_/trait_control_). The data points used for training (purple) or testing (orange) the model are shown. **(B)** Feature-scaling analysis, which normalises the independent features present in the data in a fixed range, was used to indicate the importance of the traits applied in the machine learning model in (A) that predicted the salt-induced changes in rosette area. **(C)** A second gradient-boosted tree model was developed using 50% of the phenotypic data to predict the salt-induced changes in rosette area (area_salt_/area_control_) based on all the measured traits (trait_salt_/trait_control_), with the exception of rosette perimeter and the greenness hues. **(D)** Feature-scaling analysis of the traits applied in the machine learning model in (C) that predicted the salt-induced changes in rosette area.

**Figure S8.**
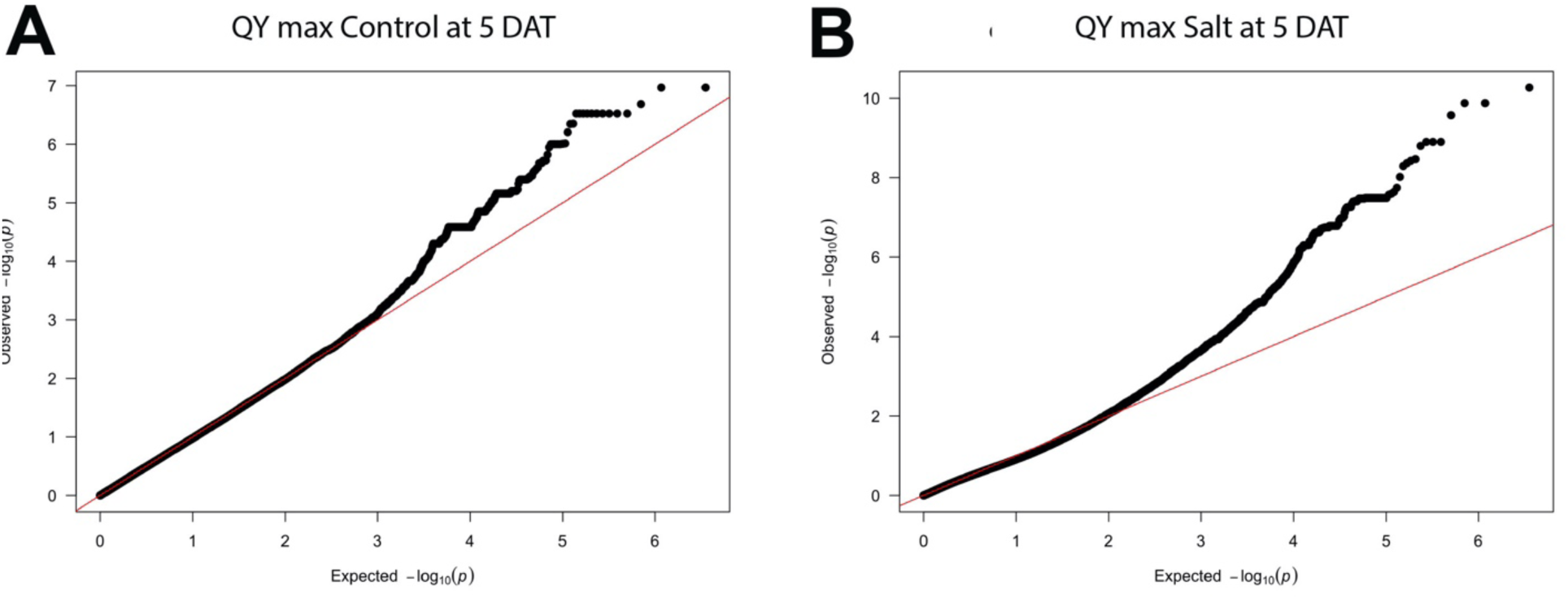
Model fitness of the genome-wide association studies (GWAS) conducted on the dark-adapted maximum quantum yield (QY max). Quantile-quantile (Q-Q) plots were generated to reflect the fitting of the single-trait GWAS models of QY max measured five days after salt treatment under **(A)** control and **(B)** salt stress conditions.

**Figure S9.**
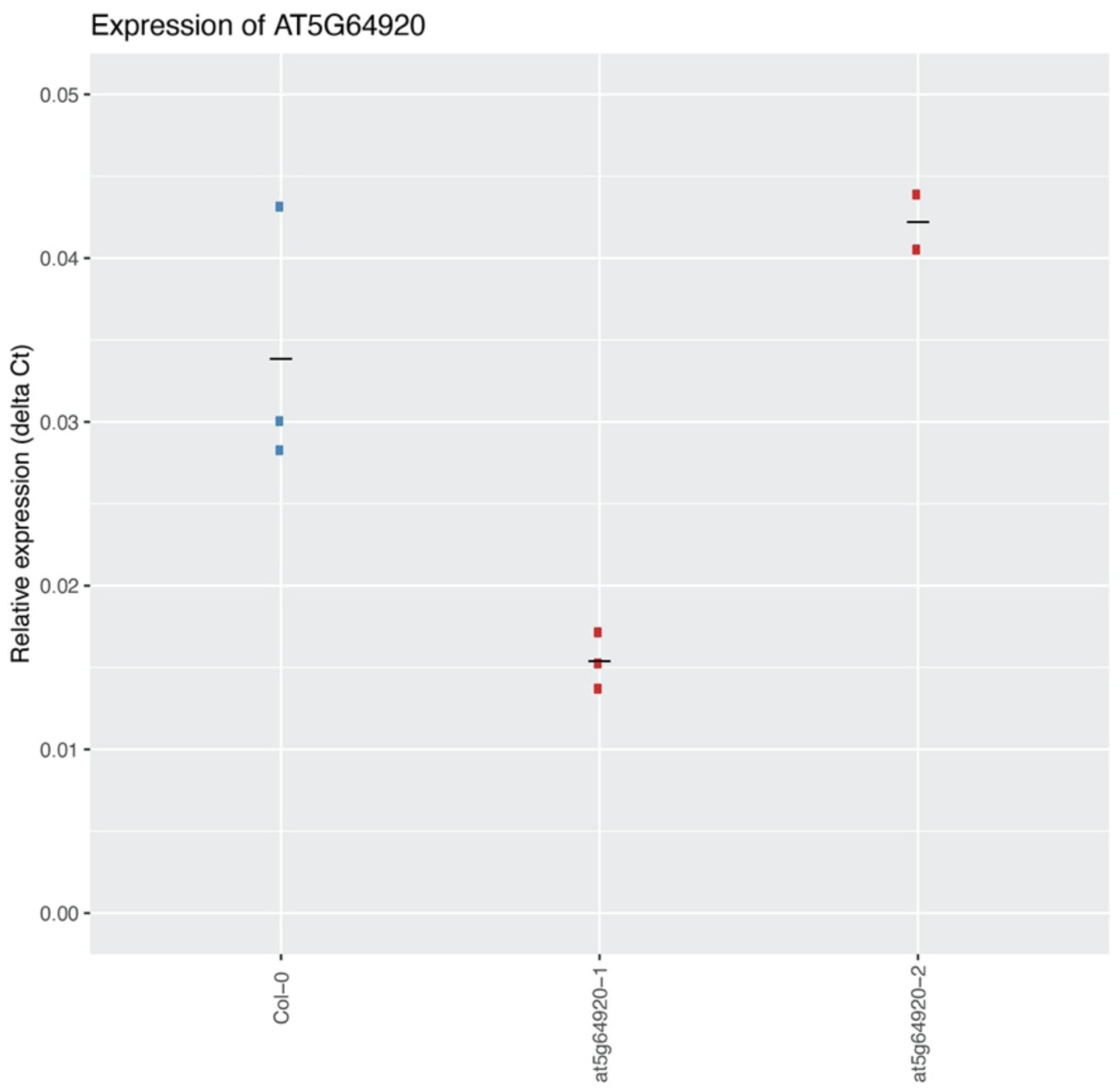
The normalised expression of the CIP8 mutant lines. The gene AT5G64920 is annotated as the constitutive photomorphogenic 1 COP1-interacting protein 8 (CIP-8). The expression of AT5G64920, encoding CIP8, was examined in Col-0 and two T-DNA insertion lines, *at5g64920-1* (SALK_138967) and *at5g64920-2* (SALK_023424), in the rosettes of three-week-old plants grown under control conditions. The expression was normalised using the delta-Ct method with the expression of actin as an internal control. Each point represents the normalised expression of three biological replicates, while the horizontal lines represent the mean expression value of each genotype.

**Figure S10.**
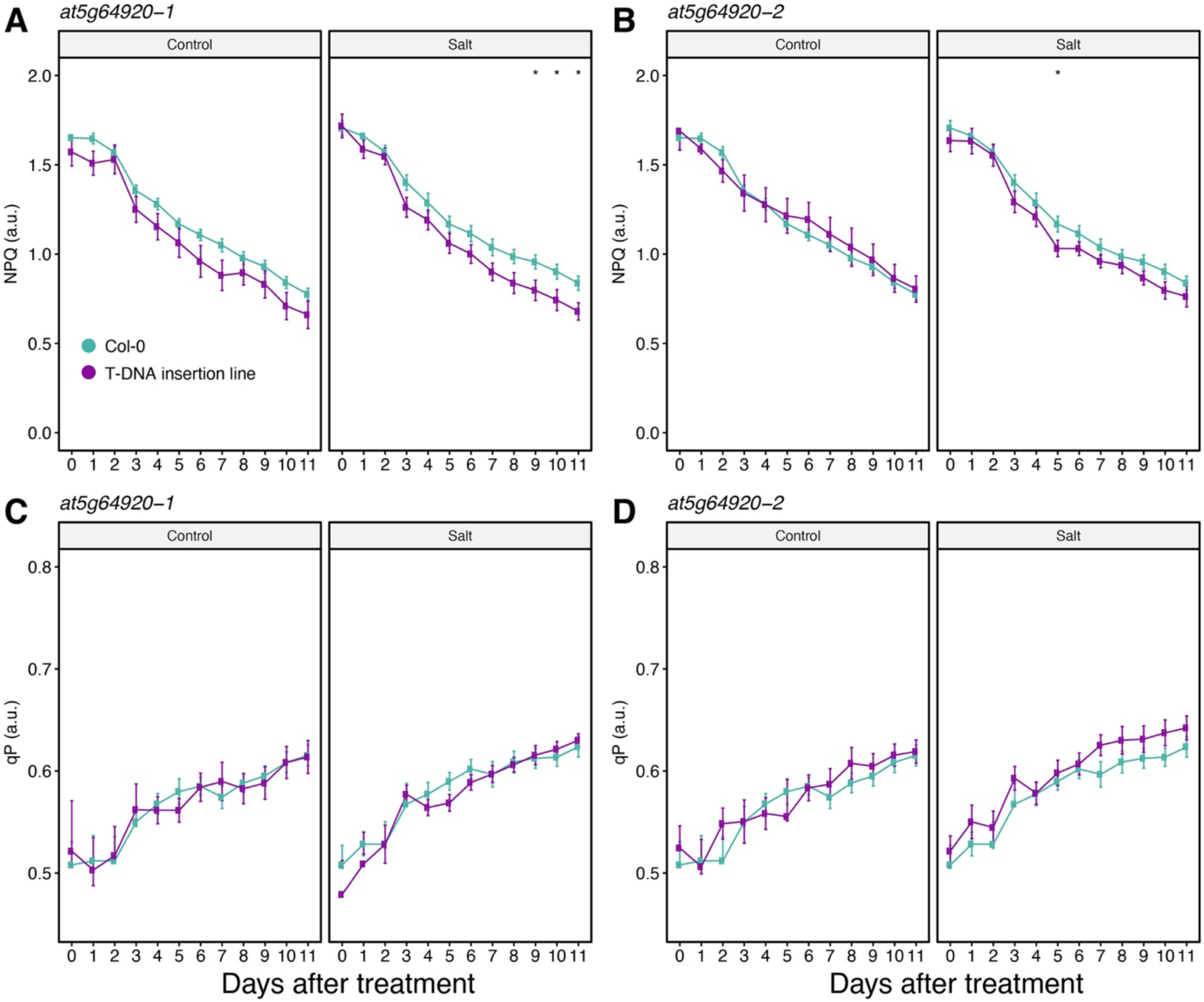
Additional phenotypes of CIP8 mutant lines. The gene AT5G64920 is annotated as the constitutive photomorphogenic 1 COP1-interacting protein 8 (CIP-8) and the salt stress responses of the two mutant alleles of AT5G64920; *at5g64920-1* (SALK_138967) and *at5g64920-2* (SALK_023424) were examined for 11 consecutive days using high-throughput phenotyping. The two-week-old plants were treated with control conditions or a final concentration of 100 mM NaCl. **(A,B)** Non-photochemical quenching [NPQ=(F_m_–F_m_′)/F_m_′)] as well as **(C,D)** photochemical quenching [qP=(F_m_′–F_t_)/(F_m_′–F_o_′)] of the background Col-0 line (blue) and the mutants *at5g64920-1* and *at5g64920-2* (purple) were measured. The asterisks indicate significant differences between Col-0 and the T-DNA insertion line, where *, **, *** and **** indicate *p*-values < 0.05, 0.01, 0.001 and 0.0001, respectively, as determined by one-way ANOVA. The lines represent the mean values calculated per genotype and condition (n=12), while the error bars indicate the standard error of the mean.

**Figure S11.**
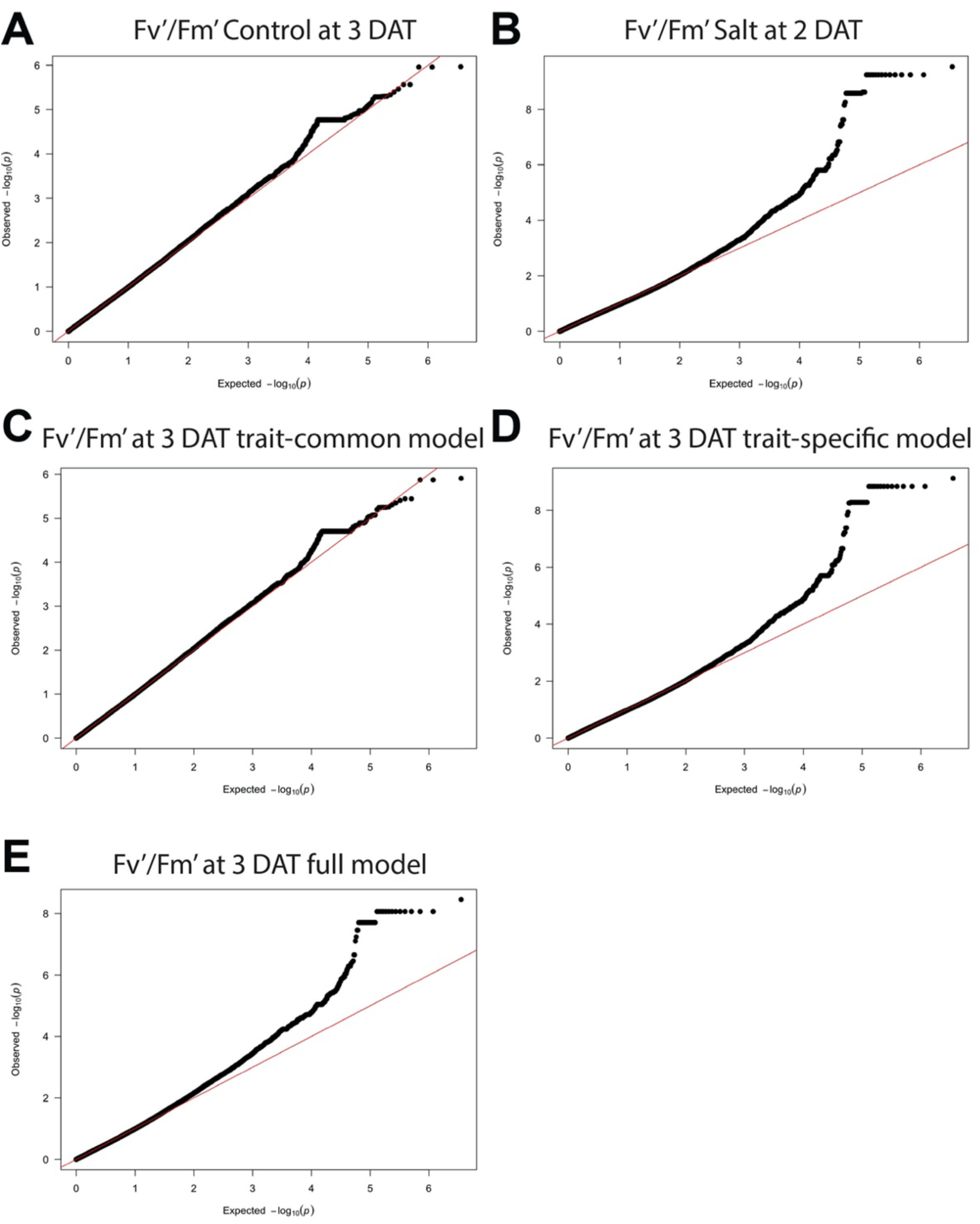
Model fitness of the genome-wide association studies (GWAS) conducted on the light-adapted quantum yield (F_v_′/F_m_′). Quantile-quantile (Q-Q) plots were generated to reflect the single- and multi-trait mixed models (MTMM) GWAS model fitting of the light-adapted quantum yield (F_v_′/F_m_′) measured three days after salt treatment using the highest photon irradiance (440 μmol m^-2^ s^-1^) in the light curve protocol of chlorophyll fluorescence imaging. The Q-Q plots were generated for associations under **(A)** control and **(B)** salt stress conditions, in addition to **(C)** the interaction between genotype and the environment (full test) as well as **(D)** the environment (trait-specific) and **(E)** genotype (trait-common) tests.

**Figure S12.**
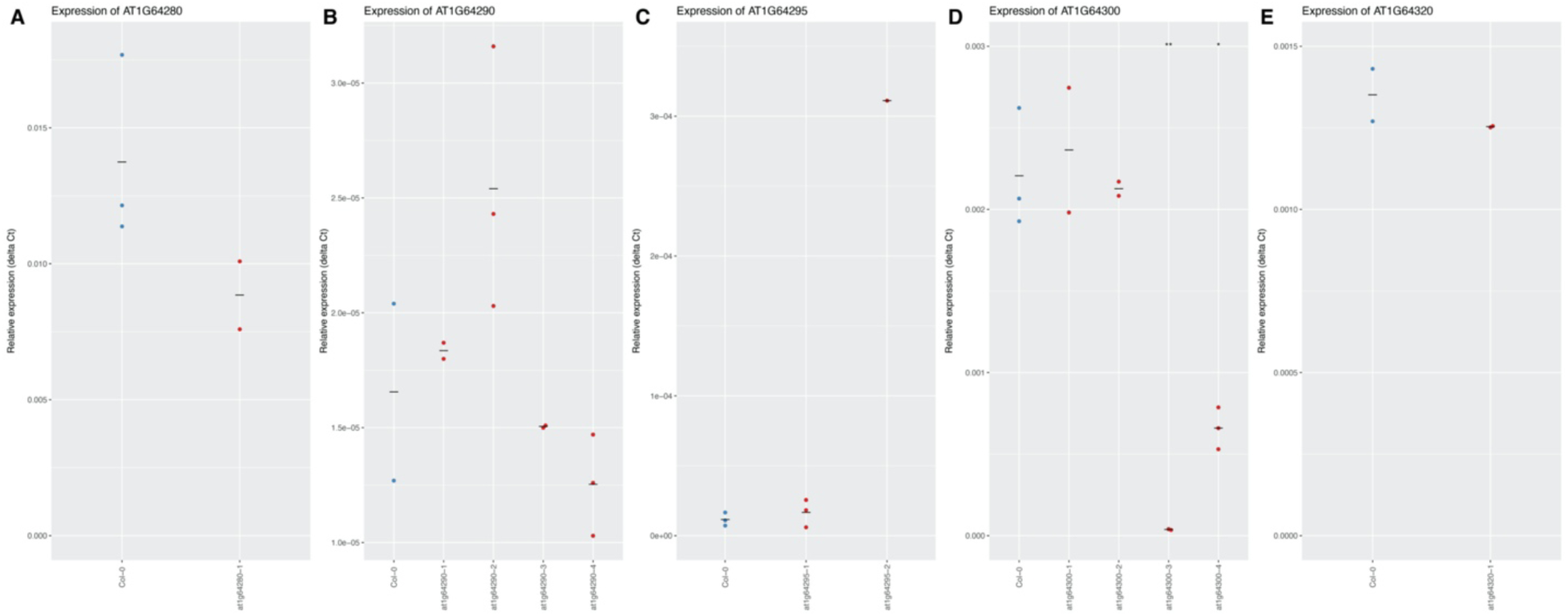
The normalised expression of the mutant lines within the F_v_′/F_m_′ locus. The gene expression of **(A)** AT1G64280, **(B)** AT1G64290, **(C)** AT1G64295, **(D)** AT1G64300 and **(E)** AT1G64320 was examined in Col-0 and T-DNA insertion lines, which included *at1g64280-1* (SALK_012492), *at1g64290-1* (SALK_000781)*, at1g64290-2* (SALK_084758)*, at1g64290-3* (SALK_011135)*, at1g64290-4* (SALK_114448), *at1g64295-1* (SALK_112855)*, at1g64295-2* (SALK_017248)*, at1g64300-1* (SALK_042948)*, at1g64300-2* (SALK_096562)*, at1g64300-3* (SALK_041569)*, at1g64300-4* (SALK_063928) and *at1g64320-1* (SAIL_1247_G09), in the rosettes of three-week-old plants grown under control conditions. The expression was normalised using the delta-Ct method with the expression of actin as an internal control. Each point represents the normalised expression of three biological replicates, while the horizontal lines represent the mean expression values per genotype. The asterisks indicate significant differences between Col-0 and the T-DNA insertion line, where *, **, *** and **** indicate *p*-values < 0.05, 0.01, 0.001 and 0.0001, respectively, as determined by a t-test.

**Figure S13.**
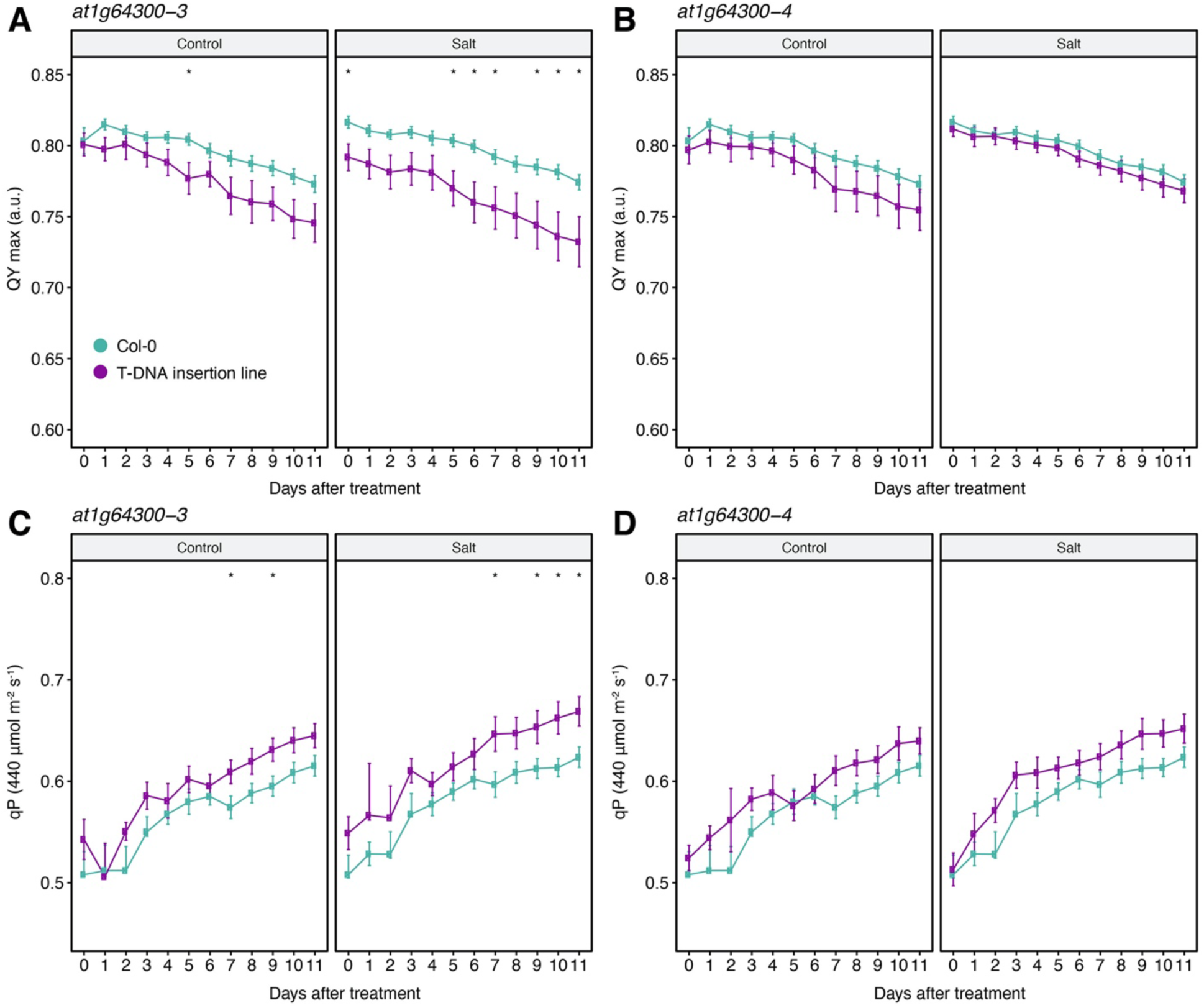
Additional mutant traits of the salt-affected photosynthetic efficiency (SAPE) gene. The gene AT1G64300 is annotated as a protein kinase and the salt stress responses of the two mutant alleles of AT1G64300, i.e. *at1g64300-3* (SALK_041569) and *at1g64300-4* (SALK_063928), were examined for 11 consecutive days using high-throughput phenotyping. The two-week-old plants were treated with control conditions or a final concentration of 100 mM NaCl. **(A,B)** Maximum quantum yield (QY max) in the dark-adapted state and **(C,D)** photochemical quenching [qP=(F_m_′–F_t_)/(F_m_′–F_o_′)] of the background Col-0 line (blue) and the mutants *at1g64300-3* and *at1g64300-4* (purple) were measured. The asterisks indicate significant differences between Col-0 and the T-DNA insertion mutant, where *, **, *** and **** represent the *p*-value < 0.05, 0.01, 0.001 and 0.0001, respectively, as determined by one-way ANOVA. The lines represent the mean values per genotype and condition (n=12), while the error bars indicate the standard error of the mean.

**Figure S14.**
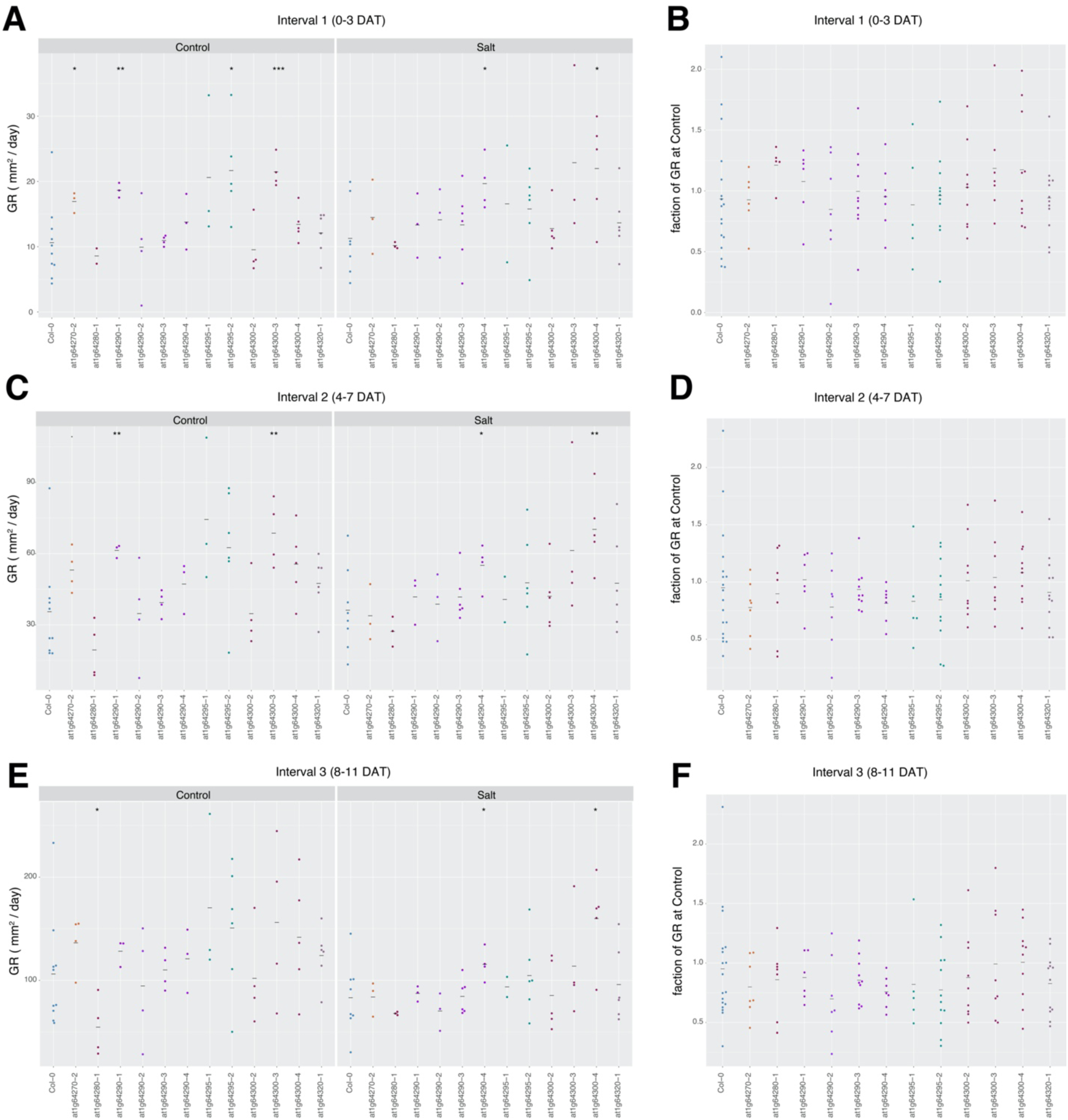
The calculated growth rates and salt tolerance indices of the T-DNA insertion lines within the F_v_′/F_m_′ locus. The salt stress responses of Col-0 and T-DNA insertion lines, which included *at1g64280-1* (SALK_012492), *at1g64290-1* (SALK_000781)*, at1g64290-2* (SALK_084758)*, at1g64290-3* (SALK_011135)*, at1g64290-4* (SALK_114448), *at1g64295-1* (SALK_112855)*, at1g64295-2* (SALK_017248)*, at1g64300-1* (SALK_042948)*, at1g64300-2* (SALK_096562)*, at1g64300-3* (SALK_041569)*, at1g64300-4* (SALK_063928) and *at1g64320-1* (SAIL_1247_G09) were examined for 11 consecutive days using high-throughput phenotyping. The two-week-old plants were treated with control conditions or a final concentration of 100 mM NaCl. The change in area through time, i.e. the growth rate, was estimated using a linear function in R during intervals 1, 2 and 3 **(A, C, E)** from 0 to 3, 4 to 7 and 8 to 11 days after treatment (DAT), respectively, which was also used to calculate the **(B, D, F)** growth maintenance by dividing the growth rate estimated under salt stress by the growth rate under control conditions per interval. Each point represents data from 12 biological replicates, while the horizontal lines represent the mean value per genotype and condition. The asterisks indicate significant differences between Col-0 and the T-DNA insertion line, where *, **, *** and **** indicate *p*-values < 0.05, 0.01, 0.001 and 0.0001, respectively, as determined by one-way ANOVA.

